# KIAA1217/SKT and its paralog p140Cap are centrosomal proteins that regulate ciliogenesis through Src family signaling pathway

**DOI:** 10.1101/2025.10.30.685566

**Authors:** Logan Greibill, Lucie Raimbourg, Hugo Siegfried, Jacky Bonaventure, Paola Defilippi, Laïla Sago, Virginie Redeker, Sandrine Etienne-Manneville, Khaled Bouhouche, Anne-Marie Tassin

## Abstract

Centrosomes organize the microtubule network along the cell cycle and drive cilia assembly in resting cells to allow sensory or motility functions. Defects in cilia formation underlie ciliopathies and skeletal disorders, but the molecular regulators that couple centrosomes to signaling and cytoskeletal networks remain incompletely defined. Through BioID screening for CYLD interactors, we identify KIAA1217/SKT as a centrosomal and microtubule plus-end protein that also associates with focal adhesions. KIAA1217 loss in RPE-1 cells impaired ciliogenesis, producing fewer and shorter cilia. Domain mapping revealed an N-terminal centrosomal targeting domain and an EB1-dependent targeting of KIAA1217 to microtubule tips via C-terminal SxIP motifs. Importantly, defects caused by loss of KIAA1217 or its paralog p140Cap were rescued by inhibiting actin polymerization or Src activity, indicating regulation of actin polymerization through Src family activity.

Together, our findings establish KIAA1217 as a positive regulator of ciliogenesis that integrates Src-dependent signaling, centrosomal architecture, and actin remodeling and may open new research avenues to understand KIAA1217-associated pathologies as vertebrate malformation and epithelia-mesenchymal transition.

Centrosome, Ciliogenesis, Actin, Src family signaling, KIAA217/SKT, p140Cap

## INTRODUCTION

Cilia are microtubule-based organelles protruding from the cell surface endowed with sensory and/or motility functions. In human, most cells bear a single immotile cilium, the primary cilium, and some cell types harbor one or multiple motile cilia. In all cases, ciliogenesis is a multistep process including basal bodies (BB) assembly, maturation and migration, transition zone formation, docking to the cell surface and further axonemal extension (Zhao et al., 2023). Defects in cilia formation or function lead to numerous genetic diseases collectively referred to ciliopathies.

Primary cilia form during the G0/G1 phase of cell cycle, during which the mother centriole is converted into BB and docks at a membrane (Zhao et al., 2023). Thanks to both microtubule motors and centrosomal actin filaments, myosinVa-associated vesicles dock to distal appendages (Wu et al., 2018). Membrane shaping proteins (EHD1, EHD3), together with SNARE and Rab proteins, mediate vesicle fusion to form the ciliary vesicle, while the centriolar capping proteins, such as CP110, are removed (Zhao et al., 2023). Accumulating evidence highlights the importance of a dynamic actin network during ciliogenesis (Kim et al., 2010, 2015; Pitaval et al., 2017a, 2010; Francis et al., 2011; Jewett et al., 2021a). Moreover, focal adhesion (FA) proteins, known to link the extracellular matrix (ECM) to the actin cytoskeleton, are also localized to the BB to tether it to the underlying actin cytoskeleton and regulate ciliogenesis (Antoniades et al., 2014). Accordingly, depletion of FA proteins disturbs ciliation in both single-ciliated (Failler et al., 2021) and multiciliated cells (Antoniades et al., 2014). Src tyrosine kinase family, localized to FAs, are key regulators of actin dynamics and cellular adhesions and have been proposed to regulate cilia formation (Drummond et al., 2018; Bershteyn et al., 2010). Indeed, the atypical protein kinase C (aPKC) and the protein Missing-in-Metastasis (MIM), recruited at the BB by the actin regulator Cdc42, regulate cilia formation by acting antagonistically on the Src signaling pathway. aPKC activates Src activity (Drummond et al., 2018), while MIM represses it (Bershteyn et al., 2010). In addition, several experimental results suggest links between Src pathway activation and the two forms of polycystic kidney diseases (PKD), autosomal dominant (ADPKD) and autosomal recessive (ARPKD) PKD, two developmental ciliopathies affecting kidney formation. Importantly: i) increased Src activity in kidneys correlates with disease progression in two rodent models of human ARPKD, a non-orthologous murine model (BPK) and an orthologous PCK rat model (PCK) (Sweeney et al., 2008); ii) In kidneys of ADPKD and ARPKD patients, activation of the Src pathway has been observed in cyst-lining renal epithelia (Talbot et al., 2014; Dafinger et al., 2020); iii) specific inhibitors of Src activity ameliorate the renal and biliary lesions characteristic of human ARPKD (Sweeney et al., 2008; Patel et al., 2009).

All these results suggest that regulation of Src signaling is critical to achieve proper ciliogenesis. However, how global and/or centrosomal actin dynamics affects cilia biogenesis through Src activity remains poorly understood.

Interestingly, the deubiquitinase tumor suppressor CYLD, which catalyzes the removal of K63 or linear ubiquitin chains, is a master regulator of the NF-kB pathway and has been shown to regulate ciliogenesis by preventing BB migration /anchoring (Eguether et al., 2014; Gomez-Ferreria et al., 2012; Yang et al., 2014). Independently, CYLD has been identified as a key member of an integrin-ILK and EGFR co-signaling pathway (Azimifar et al., 2012a). EGF stimulation resulted in a Src-dependent Tyrosine phosphorylation of CYLD, which relocalizes to dynamic actin-based dorsal ruffles, independently of its deubiquitinase activity (Azimifar et al., 2012a). Altogether these results suggest that CYLD might control ciliogenesis through interactions involving EGF, Src and actin. Therefore, to get more insights into CYLD-mediated regulations, we decided to search for proximity interactors of CYLD.

Here we report the functional analysis of one CYLD interactant, KIAA1217, also known as SKT in the mouse model. KIAA1217 shares up to 40% identity and 60% homology with p140Cap encoded by the *SrcIn1* gene (Salemme et al., 2021a). Recently SKT has been described as a relevant adaptor within the Post-Synaptic Density (PSD), demonstrating its interaction with PSD-95 and SHANK3, and providing insights into its role in synaptic signal transduction (Morellato et al., 2025). Our findings performed in Retinal Pigment Epithelium (RPE-1) cells revealed that both KIAA1217 and its paralog p140Cap control the ciliogenesis process through SRC family signaling and the actin network. Our results suggest, first, that the role of KIAA1217 in ciliogenesis could account for the syndromic lumbar disc herniation and vertebral malformations observed in *Kiaa1217* mutated patients. Second, KIAA1217 might be at a nexus linking centrosome, cilia and FA ensuring a cross talk between the centrosome/cilium and the ECM to regulate cilia homeostasis.

## RESULTS

In order to understand how CYLD regulates ciliogenesis, i.e. BB migration/anchoring, we decided to identify its proximity partners. For this purpose, Flp-In HEK293 cells expressing myc-BirA*-CYLD or myc-BirA* under the control of the tetracycline Tet-On system were generated. Total proteins from these lines, grown in biotin-containing medium, were extracted. Then, biotinylated proteins were purified on streptavidin beads and analyzed by mass spectrometry (MS). Proteins were ranked according to their fold change and p-value following a comparison between the two conditions. In the myc-BirA*-CYLD line, twenty-four proteins were identified with a fold-change >2 and a p-value< 0.05, compared to the myc-BirA* line (Fig. 1A-heatmap). Gene ontology enrichment analysis of these 24 proteins revealed that they are significantly enriched in GO terms related to cytoskeleton, centrosome, microtubules, adherens junction and ubiquitination (Fig. 1B). Several proteins such as SPATA2, SPATA2L, TRAF2, and CEP192 (see Pride database) already known as CYLD partners were found among our BioID/MS candidates, thus validating our BioID/MS screen. Moreover, in agreement with the centrosomal localization of CYLD, several centrosomal proteins such as CEP192, SAS6, HAUS3 and KIAA1217 were found in our BioID/MS analysis. Interactions between these proteins, established by STRING database, revealed that CYLD proximity-partners are interconnected (Fig. 1 C). CYLD-KIAA1217 interaction was confirmed by co-immunoprecipitation experiment using HEK293 cells expressing Flag-CYLD and KIAA1217-GFP (Fig. 1D). KIAA1217 was found previously in another BioID screen achieved in the HEK293 cell line using two centrosomal proteins (Ninein-like and centrobin) as a bait and overexpressed V5-tagged KIAA1217 fusion protein localized to the centrosome, suggesting that KIAA1217 could be enriched in the centrosome and play a role during ciliogenesis (Gupta et al., 2015). Since KIAA1217 has not been characterized functionally, we decided to perform a functional analysis focusing on the ciliogenesis process.

**Fig. 1:**
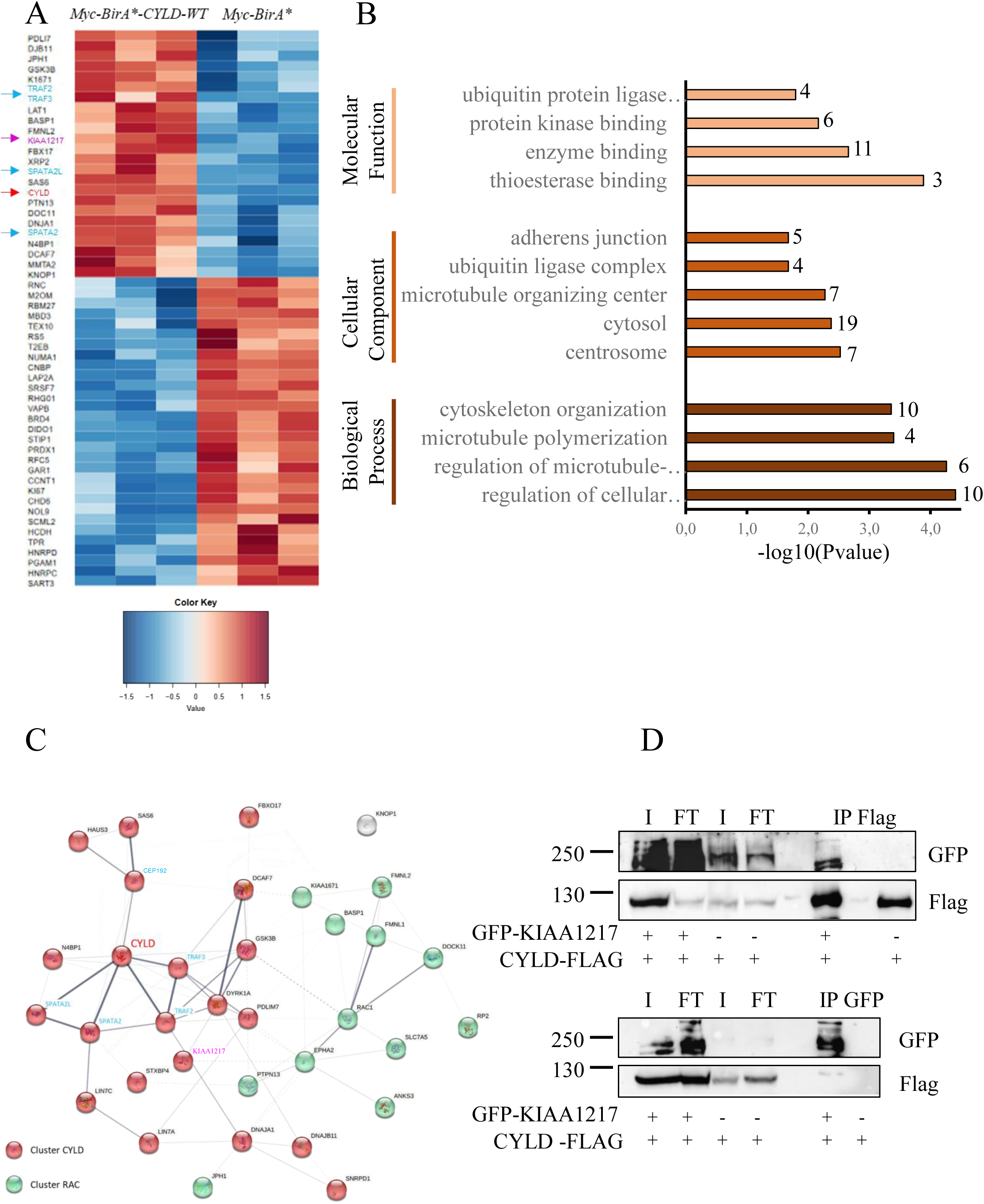
BioID/MS analysis of CYLD proximity partners identified among other KIAA1217 protein. A: Heatmap of proteins identified in myc-BirA*-CYLD + Tet versus myc-BirA*+ Tet made with Qlucore. Proteins were filtered based on a p-value < 0.05 and Fold Change > 2, intensities were log2 transformed and were displayed as colors ranging from red to blue. B: Gene ontology (GO) enrichment analysis was performed using DAVID 6.8 database. Only p-values < 0.05 and adjusted by Benjamini-Hochberg correction are displayed. The top enriched GO terms (Biological Process, Cellular Component and Molecular Function) are shown by their-log10(P-value) and the number of proteins in each set. Analysis of the 24 proteins identified by BioID/MS revealed that they are significantly enriched in GO terms related to cytoskeleton, centrosome, microtubules, adherent junctions and ubiquitination. C: The network was constructed using STRING DB representing a full STRING network at medium confidence (0.400) with disconnected nodes hidden. Lines of different thicknesses between nodes symbolize the Edge confidence from medium to the highest. Centrosomal proteins already known as CYLD interactants appear in blue. D: Co-immunoprecipitation experiment made in HEK293 cells co-expressing either GFP-KIAA1217 /CYLD-Flag or GFP/CYLD-Flag. Cell extracts were immunoprecipitated with GFP or Flag traps. The input (I), the unbound fraction (FT) as well as the bound fraction (IP) were separated on SDS-PAGE and revealed with either rabbit polyclonal anti-GFP or rabbit polyclonal anti-Flag antibodies. As expected from our BioID/MS results, CYLD-Flag immunoprecipitates GFP-KIAA1217.

### KIAA1217 and p140Cap contribute to cilia formation

Mutations in human *Kiaa1217* gene have been previously involved in various skeletal diseases,(Al Dhaheri et al., 2020), which might reflect ciliogenesis defect. These results and a high-throughput ciliogenesis screen (Gupta et al., 2015) prompted us to analyze in detail the involvement of KIAA1217 in cilia formation. KIAA1217 inactivation by two sets of siRNA (a pool of 4 siRNAs and a single one), led to a significant reduction of full-length KIAA1217 (about 80%) by western blotting (WB), thus demonstrating the efficiency of the siRNA (Fig. S1). The analysis of cilia formation 24 h after serum starvation, showed a significant reduction of the proportion cells in KIAA1217^siRNA^ condition (about 30% of ciliated cells) compared to the control^siRNA^ (70%) (Fig. 2, A-C). In addition, when primary cilia were present in KIAA1217 depleted cells, they were much shorter than in control cells (Fig. 2, A,C): after the KIAA1217 knockdown, about 70% of ciliated cells showed a cilium measuring less than 3µm while 80% of control cells display a cilium longer than 3µm. Both the significant reduction in ciliogenesis percentage and ciliary length following KIAA1217 knockdown strongly suggest that KIAA1217 is required for cilia formation and/or elongation. This defective ciliogenesis could not be attributed to an inefficient cell cycle arrest, as the majority of control and KIAA1217-depleted cells were arrested in the G1/G0 phase (Fig. S2). RPE-1 stable cell line expressing GFP-KIAA1217 construct resistant to KIAA1217^siRNAcustom4^ partially rescued the number of ciliated cells (Fig. 2F). We noticed that the GFP-KIAA1217 cell line displayed a reduced number of ciliated cells in control^siRNA^. This could be due to the overexpression of GFP-KIAA1217 or to some confluency during the establishment of the cell line. Since KIAA1217 is a paralog of the p140Cap protein, that binds and activates the C-terminal Src kinase (CSK) to inhibit Src signaling, the putative involvement of p140Cap in cilia formation was investigated. Depletion of p140Cap in RPE-1 cells using a pool of four siRNAs reduced dramatically cilia formation by 70% (Fig. 2, D-E). We verified by WB that the different siRNA we used, were indeed reducing the expression of the targeted protein (Fig. S5). Depletion of KIAA1217 by siRNA did not affect p140Cap expression and conversely depletion of p140Cap had no effect on KIAA1217 level (Fig. S5). Altogether, these results show that the depletion of either KIAA1217 or p140Cap prevent cilia formation. It should be noted that depletion of p140Cap had a stronger effect on cilia formation than KIAA1217 knock-down.

**Fig. 2:**
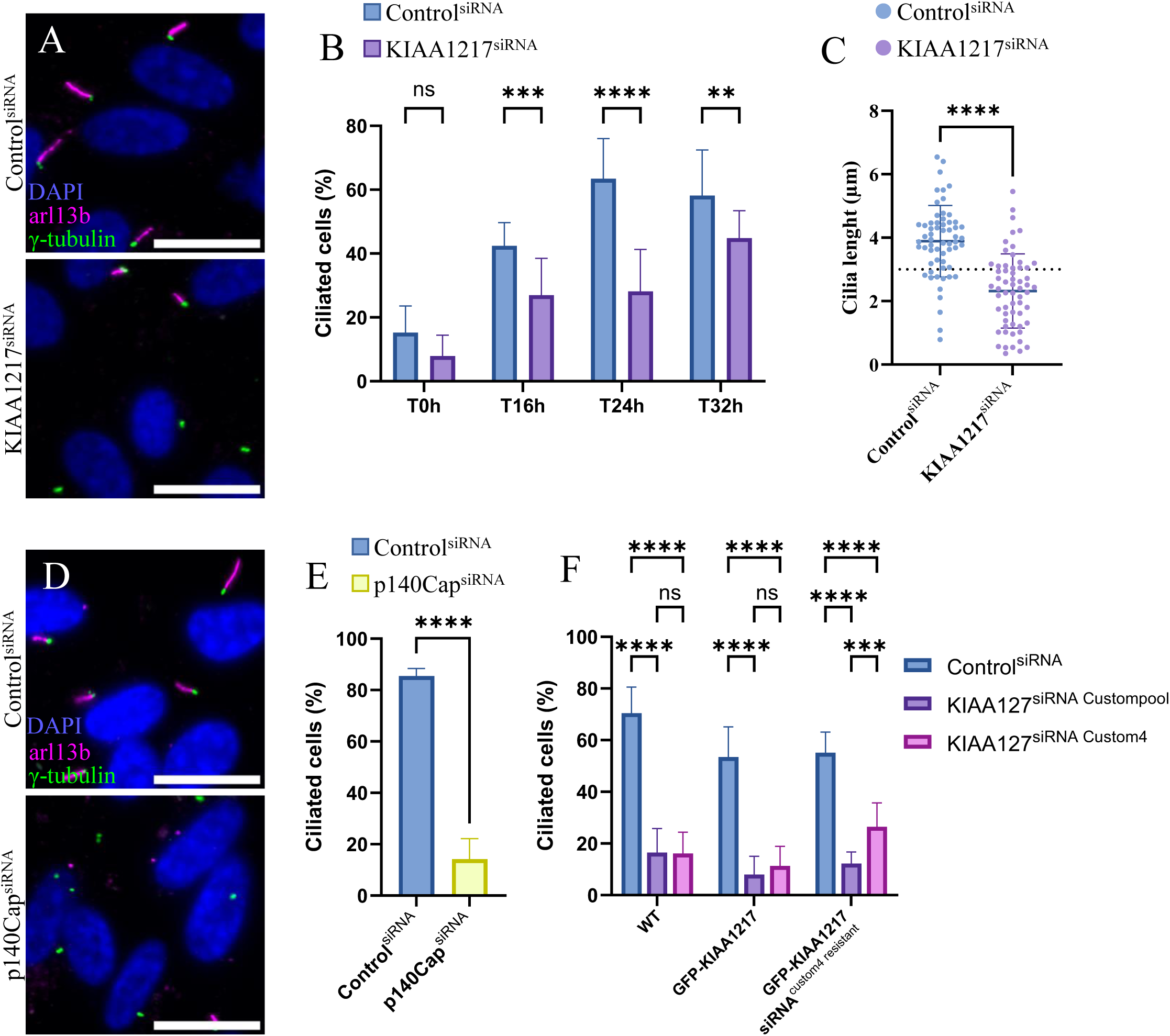
The depletion of KIAA1217 or p140Cap prevents ciliogenesis. A: RPE-1 cells were transfected with either control^siRNA^ (scrambled) or KIAA1217^siRNACustom^ ^pool^. Representative images of control or KIAA1217 depleted cells stained with both anti-arl13b (decorating the cilium) and γ-tubulin antibodies (staining the centrosome) 24 h after serum starvation. Scale bars: 20µm B: Quantification of ciliation in control and KIAA1217-depleted cells after indicated times of serum starvation. The bar graph indicates the average from two independent experiments. Cell number analyzed for each condition > 250. The bar graph represents the mean number of ciliated cells number at different times after serum starvation in control^siRNA^ and KIAA1217^siRNACustompool^. All data are presented as average ± SD. **p=0.0046 / ***p=0.0002 / ****p <0.0001 (Two-way ANOVA followed by Šídák’s multiple comparisons test). C: Quantification of ciliary length upon transfection with respective siRNA as presented in B. Dot plot represents ciliary lengths measured upon control (blue) or KIAA1217 (purple) depletion in two independent experiments. Cilia number analyzed for each condition: 60. All data are presented as average ± SD. ****p <0.0001 (Unpaired t test) D: RPE-1 cells were transfected with either control^siRNA^ or with p140Cap^siRNA^ (smartpool). Representative images of control or p140Cap depleted cells stained with both anti-arl13b and γ-tubulin antibodies 24 h after serum starvation. Scale bars: 20µm. E: Quantification of ciliation in control (black) and p140Cap-depleted cells (grey) 24 h after serum starvation. The bar graph indicates the average from two independent experiments. Total cell number analyzed for each condition >700 in 2 independent replicates. All data are presented as average ± SD. ****p <0.0001 (Unpaired t test) F: Graph showing control RPE-1 cells (WT) depleted for control^siRNA^, KIAA1217^siRNACustom^ ^pool^ and KIAA1217^siRNACustom4^ as well as RPE-1 cells stably expressing GFP-KIAA1217 and RPE-1 cells stably expressing GFP-KIAA1217^siCustom4resistant^. As expected both KIAA1217^siRNACustompool^ and KIAA1217^siRNACustom4^ decrease dramatically the number of ciliated cells. Similar results were observed for the stable GFP-KIAA1217 RPE-1 cell line. However, in GFP-KIAA1217^siCustom4^ ^resistant^ cells, KIAA1217^siRNACustom^ ^pool^ decreases the number of ciliated cells to about 10% whereas the KIAA1217^siRNACustom4^ only decreases this number to about 25%, suggesting that the decreased ciliation is partially rescued by the expression of GFP-KIAA1217^siCustom4resistant^. Total cell number analyzed for each condition >99 in 2 independent replicates. All data are presented as average ± SD. ***p<0.001 / ****p <0.0001 (Two-way ANOVA followed by Tukey multiple comparisons test)

### KIAA1217 is a core centrosomal protein

To determine the subcellular localization of KIAA1217 at the endogenous level, we performed immunofluorescence (IF) experiments using three different antibodies, two home-made and one commercial targeting 3 different regions of KIAA1217 (Fig. S1 A) (see material and method section). We first evaluated the efficiency of the three antibodies in WB and IF (Fig. 3 and Fig. S1). Three affinity-purified antibodies decorated the centrosome in U-2 OS and RPE-1 cell lines as demonstrated by co-labeling with γ-tubulin antibodies (Fig. 3A and Fig. S3 A). EVITT-DTP antibodies gave a much weaker IF signal than the two others (Fig. S3 A). The localization of KIAA1217 was confirmed in both mono and multiciliated cells, in which KIAA1217 concentrated at the BB (Fig. 3, C-D) KIAA1217 localization in RPE-1 cells was not sensitive to microtubule depolymerization (Fig. S3 D). KIAA1217 association with centrosomes was confirmed by immunofluorescence on purified centrosomes (Fig. 3E, left) and by WB analysis of centrosomal fractions compared to the soluble and insoluble fractions (Fig. 3E, right).

**Figure 3:**
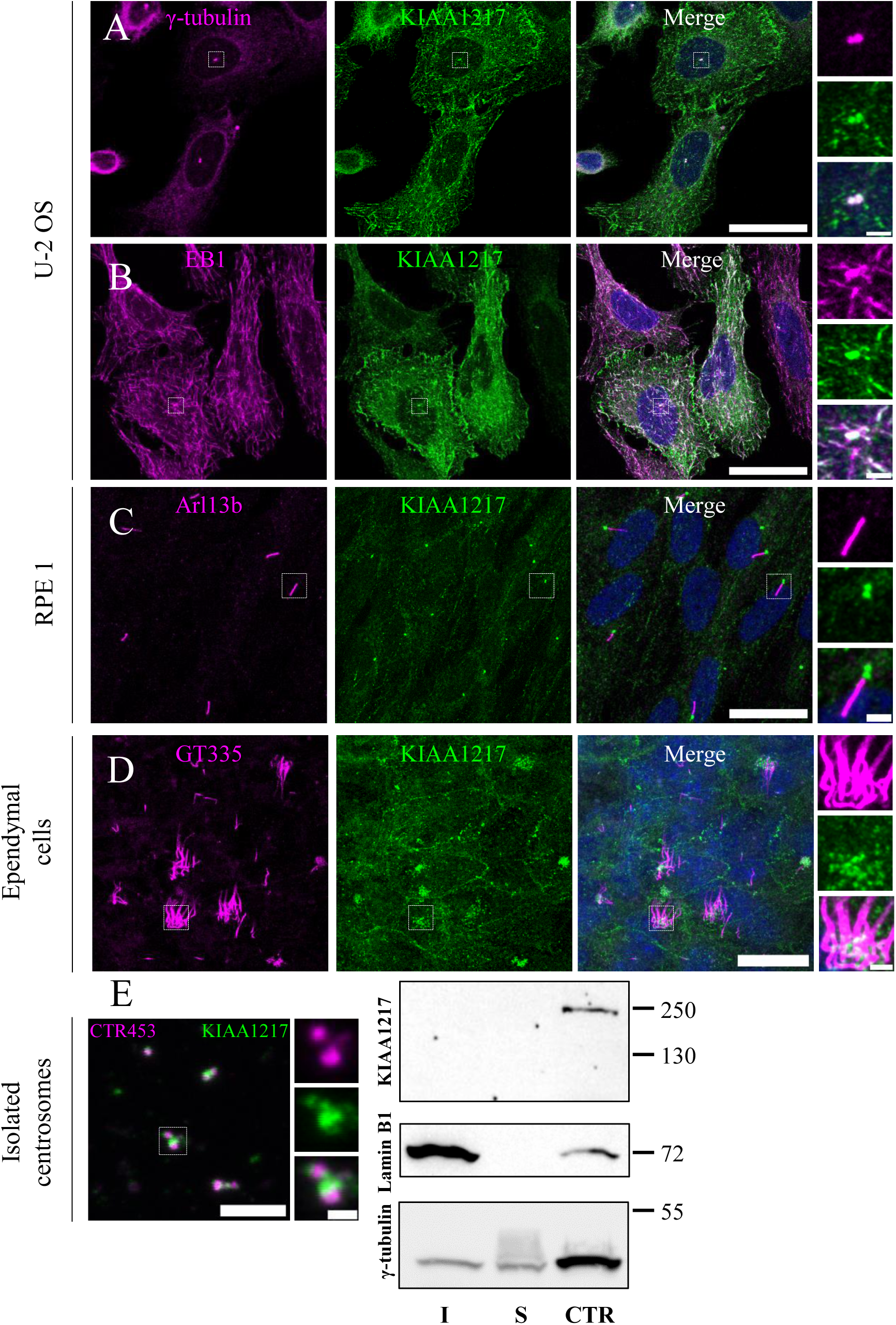
KIAA1217 is enriched at the centrosome. A-B: U-2 OS cells were stained with rabbit antibody directed against KIAA1217 (Proteintech) and the mouse gtu88 antibody recognizing γ-tubulin (A) or monoclonal EB1 antibody (B). A: The centrosome is decorated by both γ-tubulin and KIAA1217 antibodies similarly as shown on the magnification (right panel). In addition to the centrosome staining, KIAA1217 antibody recognizes numerous dots in the cytoplasm that partially colocalize with EB1 comets. Scale bars: 20µm / 2µm in magnification. C: RPE-1RPE-1 cells were starved to induce cilia formation. The primary cilium is decorated with Arl13b antibody. As expected from the centrosomal staining, KIAA1217 (Proteintech) antibody labelled the BB from which the cilium emerges. Scale bars: 20µm / 2µm in magnification. D: ependymal cells were decorated with both GT335 antibody (recognizing polyglutamylated tubulin) and KIAA1217 NKF-KPT antibody; GT335 decorates as expected cilia, and numerous dots close to the cilia are observed with KIAA1217 antibodies suggesting that in multiciliated cells KIAA1217 is located at BB. In addition, cellular junctions are also labelled. Scale bars: 20µm / 2µm in magnification. E: Centrosome purification performed on KE37 cells. In the left panel isolated centrosomes are decorated with either γ-tubulin or KIAA1217 antibodies (NKF-KPT). Scale bars: 20µm / 2µm in magnification. In the right panel, WB analysis of the triton X-100 insoluble (I) and soluble (S) fractions as well as the purified centrosome fraction revealed by KIAA1217 EVITT-DTP or γ-tubulin antibodies. Note that KIAA1217 is enriched in the centrosome fraction as well as γ-tubulin and displays one band at 250kDa, whereas Lamin is enriched in the insoluble fraction.

In addition to the centrosomal localization, KIAA1217 also appears as cytoplasmic dots and filaments (Fig. 3A-B). IF experiment using double labeling with EB1 and KIAA1217 antibodies, showed a partial co-localization, suggesting that KIAA1217 might localize at microtubule plus-end (Fig. 3B). In addition, KIAA1217 was also observed at cell-cell contacts, which agrees with the results obtained in Fülle et al. (Fülle et al., 2024) (Fig. 3, A-B, E-F).

To confirm the endogenous KIAA1217 localization, we transiently over-expressed GFP-KIAA1217 in RPE-1 and U-2 OS cell lines (Fig. 4). At the low expression level, GFP-KIAA1217 was only detected at the centrosome (Fig. 4A). At moderate expression level, KIAA1217-GFP colocalized with EB1 confirming its recruitment at the growing microtubule ends (Fig. 4B). Finally, GFP-KIAA1217 also localized at FA labeled with anti-talin and anti-paxilin antibodies (Fig. 4, C-D). At a higher expression level, GFP-KIAA1217 was shown not only at the +ends but also along microtubules (Fig. 4E). Interestingly, some stress fibers were observed when IF was performed with phalloidin and anti-GFP antibodies using glutaraldehyde-formaldehyde as a fixative (Fig. S4 A).

**Fig. 4:**
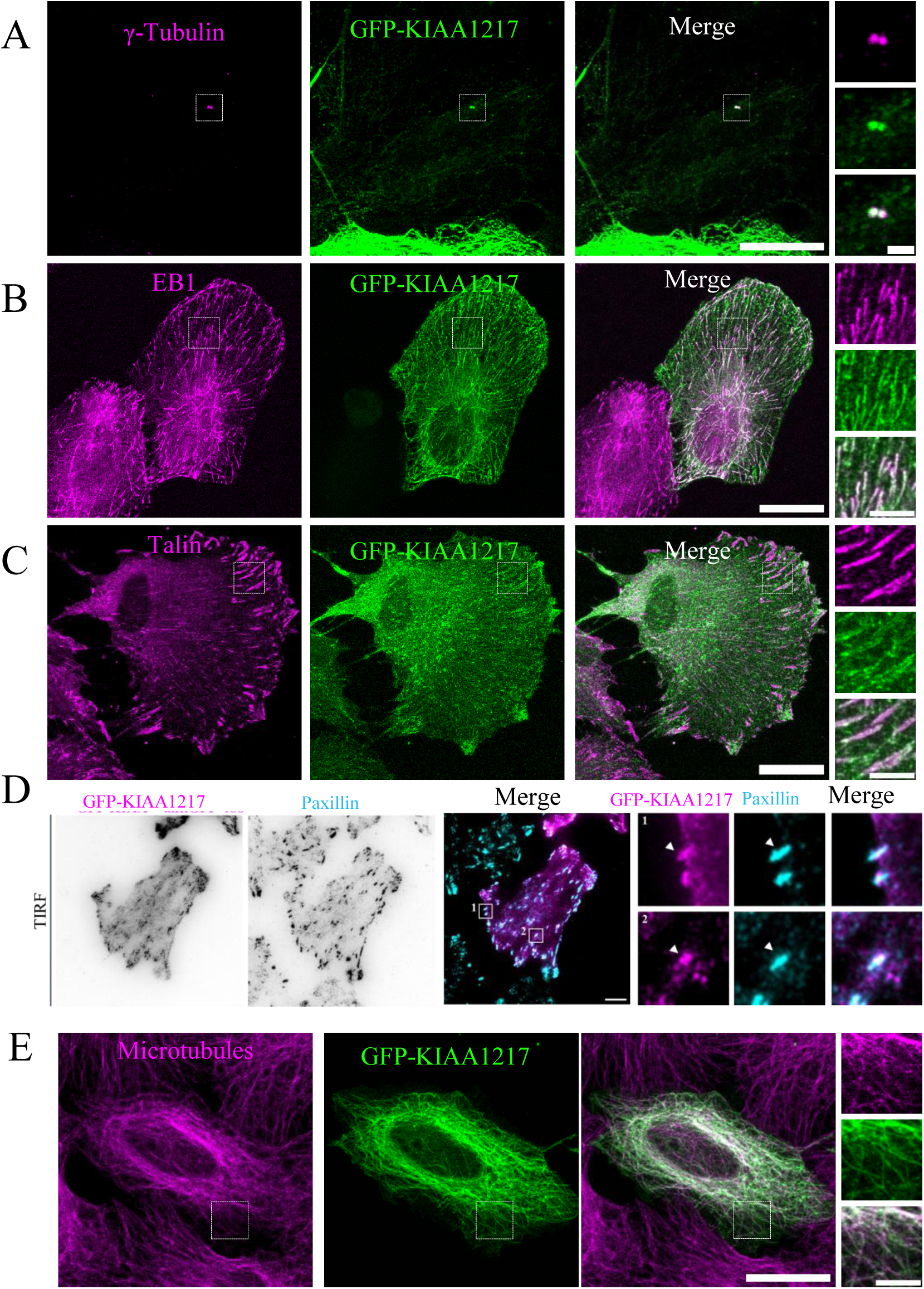
localization of GFP-KIAA1217. U-2 OS cells were transfected with GFP-KIAA1217 construct and stained by anti-GFP antibody and either anti-γ-tubulin (centrosome) (A), anti-EB1 (microtubule + Tips) (B), anti-talin (FA) (C) antibodies. Cells expressing different expression level of GFP-KIAA1217 are shown: A low expression level, B-C intermediate level. At the lowest expression level, GFP-KIAA1217 decorates only the centrosome as demonstrated with the co-labeling with anti-γ-tubulin antibody. At intermediate expression level, GFP-KIAA1217 is observed in numerous comets that colocalise with microtubule +Tips decorated with anti-EB1 antibody. Otherwise, GFP-KIAA1217 is also localized at FA as observed by the co-localization with Talin antibody (see magnification). Scale bars: 20µm / 2µm in magnification. D: Human glioblastoma cells expressing intermediate level of GFP-KIAA1217 labelled with anti-Paxillin antibody and analyzed by TIRF microscopy. Note that most of the FA are co-stained. Scale bars:10µm E: U-2 OS cells expressing GFP-KIAA1217 at the highest expression level. The microtubule network is stained by GFP-KIAA1217. Scale bars: 20µm / 2µm in magnification.

To characterize more precisely the centrosomal localization of KIAA1217, we turned to Ultrastructural Expansion Microscopy (U-ExM), a super-resolution method based on the physical expansion of the biological sample in a swellable polymer. Since the KIAA1217 antibodies we developed did not work in U-ExM, we performed this experiment on U2-OS cells expressing GFP-KIAA1217, allowing the use of GFP and tubulin antibodies. Since GFP-KIAA1217 was overexpressed, we analyzed only centrioles, which were not saturated by GFP staining. KIAA1217 formed a cylindrical cap on the outer surface at the proximal part of the two mature centrioles. Image analyses of the distribution of KIAA1217 structure relative to the centriolar wall displayed an average height of 290 nm, centered on the centriole proximal end, reminding the localization of PCM protein in U-ExM(Schweizer et al., 2021) (Fig.5 A). Comparison of cells co-stained for distal appendages marker CEP164 and KIAA1217 (Fig. S3 E) with cells labelled with either CEP164 (Fig.5B) or KIAA1217 suggested that KIAA1217 distal localization is visible only on the mother. Top-viewed centrioles showed that this proximal staining was organized in a nine-fold symmetry close to the microtubule wall at about 100 nm from the centriolar wall (Fig. 5, A,C,D). An internal staining is also observed proximally. In addition to this proximal staining, a GFP labeling was also observed at the level of the distal end of one centriole, emerging perpendicularly from the centriole wall, resembling the distal or subdistal appendages (Fig. 5, A, E). Interestingly, KIAA1217 distal staining is located at about 382nm from the proximal end whereas CEP164 is found at 464 nm. The measurements of both KIAA1217 and CEP164 staining relative to the tubulin maximal intensity signal on top viewed centrioles showed that KIAA1217 is further away to the microtubule wall than CEP164 (average distance between KIAA1217 and tubulin about 130nm, while CEP164 is about 88 nm) (Fig. 5, A-D).

**Fig. 5:**
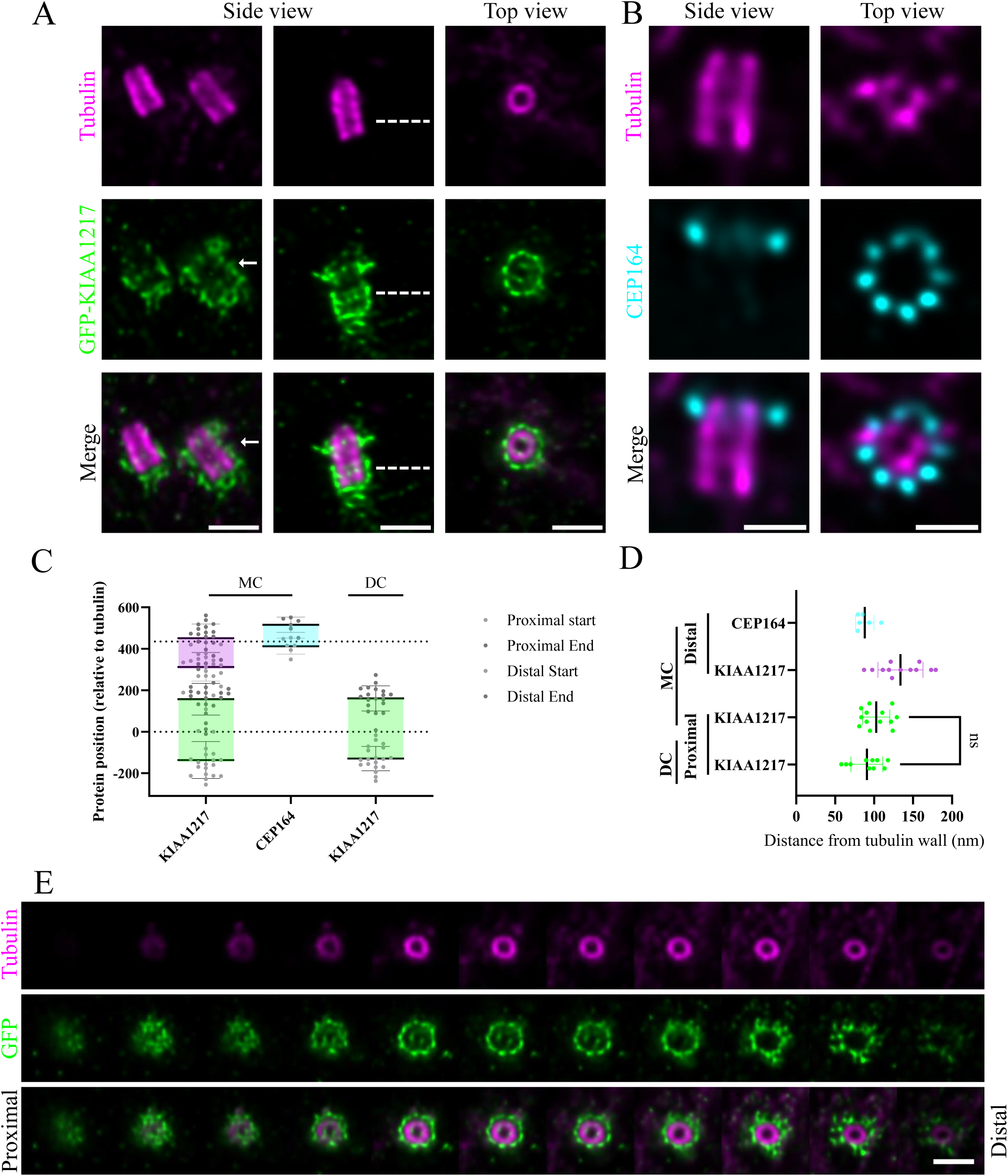
Localization of GFP-KIAA1217 by U-ExM. A: U-2 OS cells with low expression level of GFP-KIAA1217 were expanded using the U-ExM technique and stained for GFP and tubulin antibodies. The left panel shows a side view of a centrosome with a mother (arrow) and a daughter centriole, the middle panel displays the side view of a mother centriole. Mother and daughter centrioles display GFP-KIAA1217 labeling at the proximal part of the centriole; The staining embeds the proximal part of both centrioles. At the distal end, the staining is observed only on the mother centriole subdistally localized. The right panel displays a centriole observed on its top view Note that a nine-fold symmetry is observed. Scale bar: 250nm B: CEP 164 staining of the mother centriole observed by U-ExM side view (left) and top view (right). Scale bar: 250nm C: Distance between the proximal part of centriole (tubulin) and the fluorescence center of mass of GFP-KIAA1217 (green for proximal staining, brown for the distal one), CEP164 (blue) for mother and daughter centrioles. Note that the proximal staining of GFP-KIAA1217 is below CEP164 suggesting that KIAA1217 localizes to the sub-distal appendages; Error bars indicate SD. N= 20 for KIAA1217 and 5 for Cep164 D: Localization of GFP-KIAA1217 proximal (green) and distal staining (brown) and CEP164 (blue) with respect to the microtubule wall (radial position) of both the mother and daughter centrioles. Error bars indicate SD. N= 12 KIAA1217 - and 6 CEP164 mother centrioles, 10 daughter centrioles. E: Montage through the z-axis of a top view mature centriole stained for tubulin (magenta) and GFP (green). The merge channel shows that GFP-KIAA1217 is found visible the tubulin staining. This staining is detected as a ring around the proximal region of mature centrioles as soon as the tubulin signal appears. At the centriole distal part, GFP-KIAA1217 staining still illustrate a ring close to the centriolar microtubule wall, from which filaments seem to emerge. Scale bar: 500nm.

### KIAA1217 displays a centrosomal and a microtubule +Tips targeting domains

To determine which KIAA1217 domains mediate its various subcellular localizations, several truncation mutants were generated from a GFP tagged KIAA1217 version (see diagram on Fig. 6A). A cytoplasmic localization was observed for the N-terminal domain spanning the AIP3 (actin interacting protein3 domain) (AA 1-269) (Fig. S4, B,C). With the AA 1-555 construct, the centrosome was detected on a strong cytoplasmic background after methanol fixation (Fig. 6B, Fig. S4 D). An extension of the latter construct AA 1-1159 allowed the detection of GFP-positive filaments within the cytoplasm (Fig. 6C) together with the centrosome. To determine the nature of these filaments, cells were extracted with triton-X100 before methanol fixation. Under these conditions, the microtubule network was visible together with the centrosomal staining, as evidenced by the colocalization of anti-GFP and anti-tubulin stainings (Fig. 6D). The domain encompassing AA 555-1943 localizes at the centrosome with numerous dots that colocalise with microtubule +Tips (Fig 6E and Fig. S4, E,F). Triton-X100-extracted cells expressing this domain display FA localization as shown by double labeling with Talin (Fig. S4 G). Finally, microtubule +Tips were clearly distinguished on a cytoplasmic background in cells expressing KIAA1217 C-terminal domain (AA 1159-1943), as confirmed with the EB1 staining (Fig. 6F). As expected both EB1 and GFP staining completely disappeared after triton-X-100 extraction, (Fig. S4 H). These results suggest that KIAA1217 targets the centrosome by an N-terminal domain and interacts with microtubule plus-ends through its carboxy-terminal region (Fig. 6G).

**Fig. 6:**
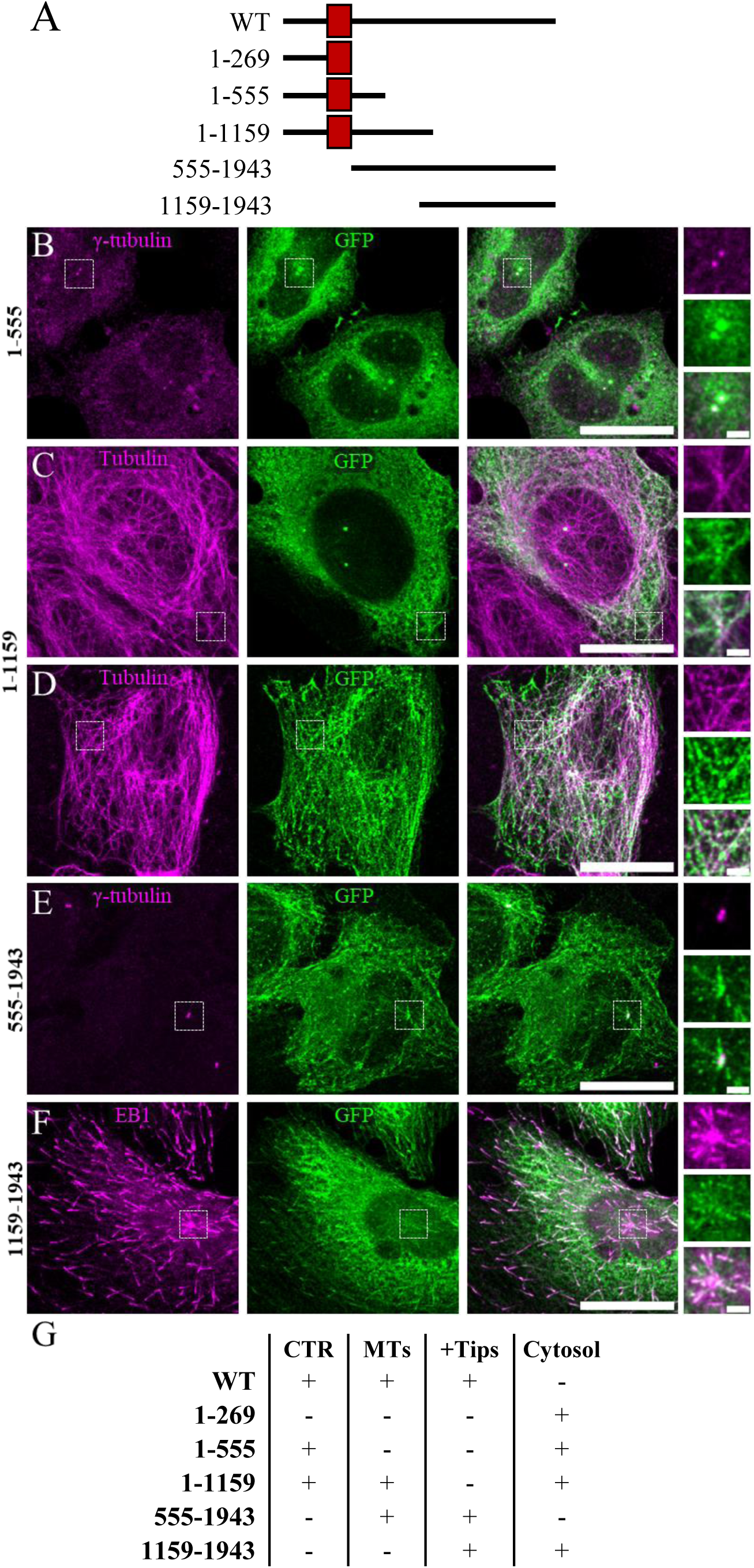
expression of constructs producing truncated KIAA1217. A: Diagram of the different GFP-KIAA1217 truncations realized. The red box indicates the AIP3 domain. B-F: U-2 OS cells expressing the GFP-tagged construct encompassing the AA 1-555 part of KIAA1217 (B), AA 1-1159 (C-D), AA 555-1943 (E), AA 1159-1943 (F). Cells were fixed with Methanol-20°C for 6 min (A, B, D, E) or fixed with Methanol-20°C for 6 minutes after 1 min permeabilization in PHEM buffer containing 0.5%triton X-100 (D). Cells were then stained with anti-GFP antibody as well as with the indicated antibodies. Scale bars: 20µm / 2µm in magnification. B: U-2 OS cells were fixed and labelled with anti-GFP and anti-γ-tubulin antibodies. The GFP staining is observed at the centrosome (arrow) as revealed by an anti-γ-tubulin antibody over a cytoplasmic staining. C: U-2 OS cells were fixed and labelled with anti-GFP and anti-α-tubulin antibodies. GFP staining decorates the centrosome (arrow). In addition, some filaments could be observed in the cytoplasmic staining. D: U-2 OS cells were fixed with methanol after tritonX-100 extraction and labelled with anti-GFP and anti-α-tubulin antibodies. TritonX-100 removed most of the cytoplasmic background and microtubules are observed as shown with-α-tubulin antibody staining. E: U-2 OS cells were fixed with methanol-20 and labelled with anti-GFP and anti-γ-tubulin antibodies. GFP decorates the centrosome (arrow) co-stained by γ-tubulin antibody, as well as comet-like emerging from the centrosome. F: U-2 OS cells were fixed and labelled with anti-GFP and anti-EB1 antibodies. With this construct the centrosome is no longer labelled with GFP antibody but is decorated with EB1 antibody; EB1 comets emerge from this dot and are co-labelled with anti-GFP antibody. A cytoplasmic staining is also observed with GFP antibody. G: Table summarizing the localizations of the different GFP-KIAA1217 truncations used: centrosome (CTR), microtubules (MTs), microtubule +Tips (+Tips) or cytoplasm (cytosol).

### KIAA1217 binds to EB1 through SxIP/SxLP motifs

To understand how KIAA1217 is targeted to the microtubule +Tips, we searched for the consensus EB1 binding motifs SxIP/SxLP in the KIAA1217 primary sequence and found eight motifs all along the protein sequence (Fig. 7A), as reported by (Jiang et al., 2012).

**FIG. 7:**
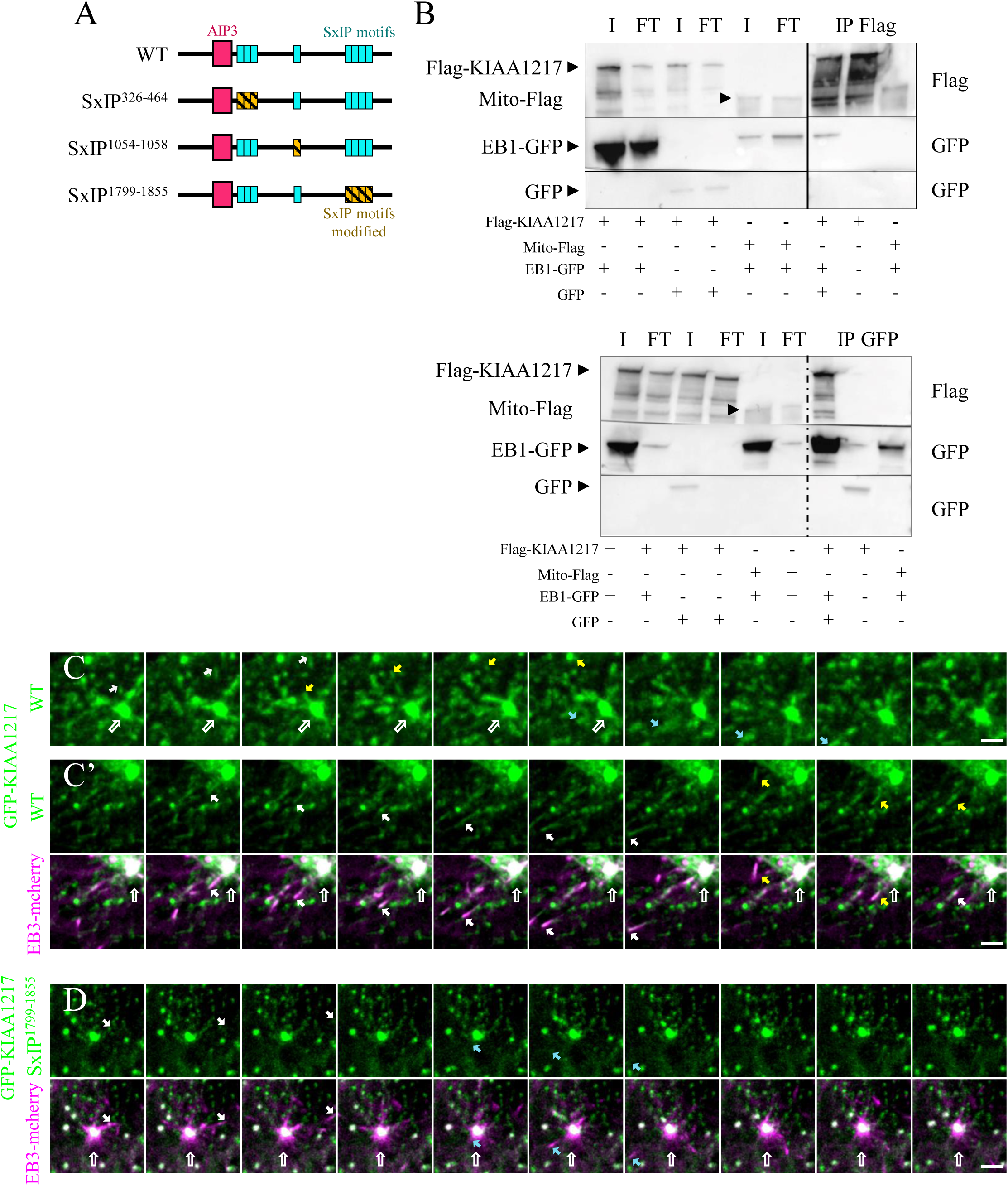
Localization of GFP-KIAA1217 regarding the microtubule +Tips protein (EB1 or EB3) A: Diagram of different GFP-KIAA1217 constructs. The red box represents the AIP3 domain, the blue boxes mark the SxIP/SxLP motifs found in the sequence. The SxIP/SxLP have been mutated into SxNN to prevent the binding to EB1 (yellow striped boxes). B: Co-immunoprecipitation experiment made in HEK293 cells co-expressing either Flag-KIAA1217/EB1-GFP, Flag-KIAA1217/GFP or mito-Flag/GFP. Cell extracts were immunoprecipitated with GFP or Flag traps. The input (I), the unbound fraction (FT) as well as the bound fraction (IP) were separated on SD-PAGE and revealed with either rabbit polyclonal anti-GFP or rabbit polyclonal anti-Flag. Note that Flag-KIAA1217 immunoprecipitates EB1-GFP but not GFP. Similarly, EB1-GFP immunoprecipitates Flag-KIAA1217 and very poorly mito-Flag. C: Time-lapse experiment: Series of 10 images were acquired every 5 seconds of RPE-1RPE-1 cells expressing GFP-KIAA1217. A comet-like structure revealed by GFP auto-fluorescence emerging from the centrosome is followed through the movie. See arrows in white, green and blue, which represent 3 distinct comets emerging from the centrosome. C’ same experimental conditions but RPE-1 cells have been co-transfected with both GFP-KIAA1217 (green) and EB3-mcherry (magenta). Two comets stained for mcherry or GFP emerge from the centrosome are pointed by a white or a yellow arrow. The centrosome is indicated by an empty arrow. Scale bars: 2µm D: Time-lapse experiment: RPE-1 cells co-expressing both GFP-KIAA1217 mutated in the four SxIP/SxLP motifspresent at the C-terminal of the protein and EB3 mCherry. Note that despite EB3-mcherry comets are observed (white arrow or blue arrow) no comets are detected with mutated GFP-KIAA1217. Scale bars: 2µm

The co-expression of Flag-KIAA1217 together with EB1-GFP in HEK293 cells, followed by IP/WB, revealed a strong interaction between both proteins (Fig. 7B). Live imaging of U-2 OS cells transiently expressing GFP-KIAA1217 (Fig. 7C and supplementary movies) showed dynamic comet-like accumulations as previously reported for EB1 and EB3, two EB family members. This result was confirmed using U-2 OS cells expressing both GFP-KIAA1217 and EB3-mcherry (Fig. 7C’, Supplementary movie1). Altogether, these results demonstrated that GFP-KIA1217 emerges from the centrosome as dynamic comets that localize with EB3-mcherry comets, in agreement with the IF data (Fig. 4B).

Finally, to demonstrate that GFP-KIAA1217 interacts with the microtubule +Tip through EB1, we generated three plasmid constructs of *GFP-Kiaa1217*, each containing some point mutations converting SxIP/SxLP motifs into SxNN motifs, that prevent EB1 binding as previously reported(Jiang et al., 2012). The three constructs produced respectively mutated SxIP/SxLP motifs either within the N-terminus of GFP-KIAA1217, or within the middle domain or within the C-terminus as shown on Fig. 7A. These constructs were transiently co-transfected with EB3-mcherry in U-2 OS cells and the dynamics of the wild type and the different mutated GFP-KIAA1217 versions were analyzed by live-imaging. As shown on Fig. 7D, the C-terminal mutated SxIP (SxIP^1799-1855^) no longer showed these dynamic comet-like structures (Supplementary movie2), whereas comets were preserved for constructs with mutant SxIP motifs within the N-terminus or middle portion. (Fig. S4, Supplementary movie3,4). Therefore, we propose that KIAA1217 binds to EB1 through the SxIP motifs localized at the C-terminus. This result is in accordance with the sub-cellular localization to the microtubule +Tips of the domain encompassing AA 1159-1943.

### p140Cap localizes to the centrosome

To understand how p140Cap control ciliogenesis, we analyzed its localization by IF in RPE-1 cells at endogenous and under overexpression of p140Cap-Flag. As shown on Fig. 8, A,B, endogenous and overexpressed p140Cap showed a centrosomal localization. Hence, the localization of both KIAA1217 and p140Cap at the centrosome suggests a potential functional redundancy between the two proteins. Therefore, we wondered whether the two proteins might interact together. IP experiments in HEK293 cells transiently overexpressing GFP-KIAA1217 and p140-Flag using both GFP and Flag Trap beads showed that GFP-KIAA1217 immunoprecipitates p140Cap-Flag and reciprocally (Fig. 8C). These results suggest that KIAA1217 and p140Cap interact and may share functions, while each protein may exert its own specific function.

**Fig. 8:**
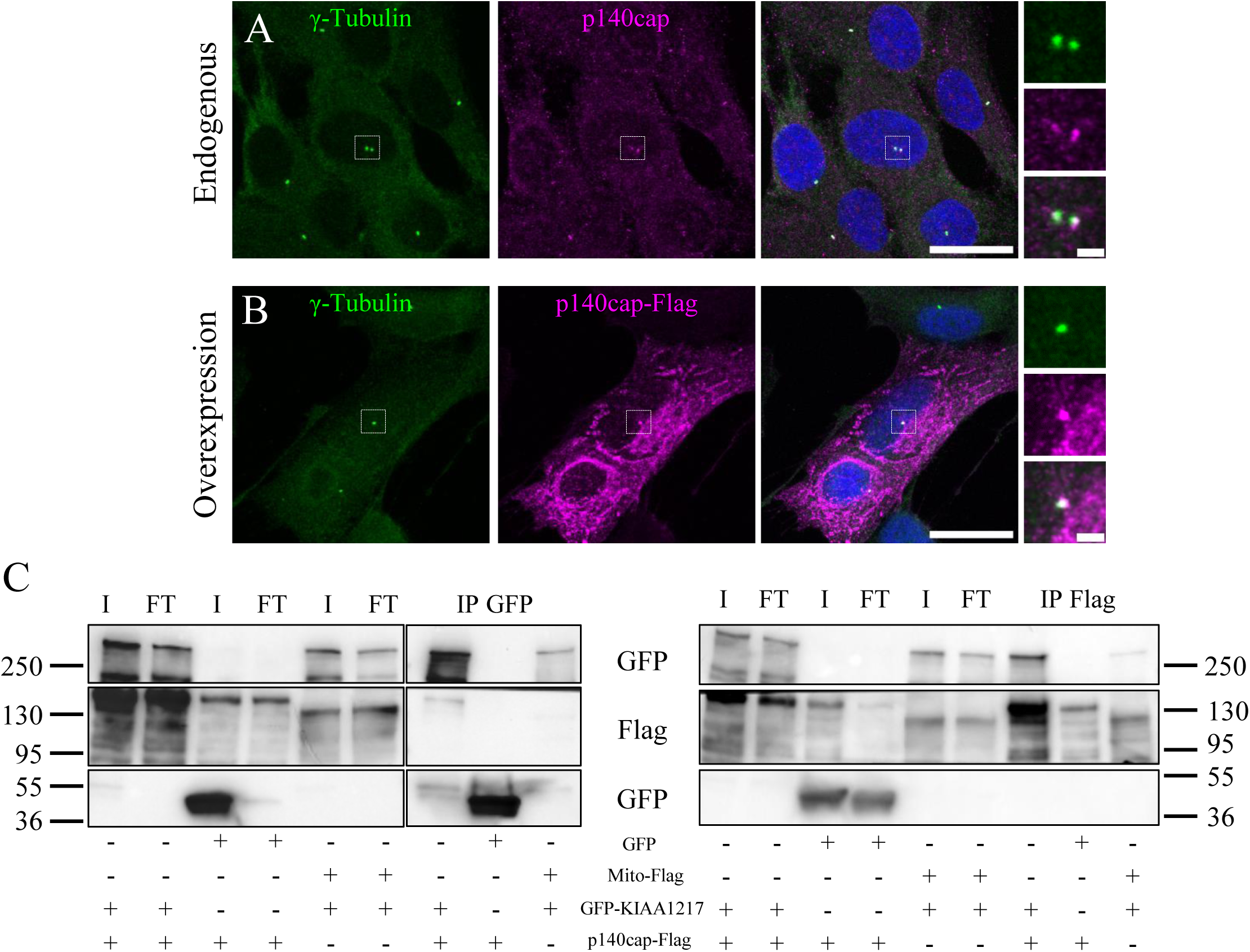
p140Cap localization in cells. A: RPE-1 cells were fixed and stained with both anti-γ-tubulin (green) and p140Cap (magenta) antibodies. P140Cap antibody decorates the centrosome as shown by the colocalization with γ-tubulin antibody. Scale bars: 20µm / 2µm in magnification. B: RPE-1 cells transfected with p140Cap-Flag were fixed and stained with both anti-γ-tubulin (gtu88, green) and rabbit anti-Flag (Magenta) antibodies. Flag antibody decorated the centrosome as expected from the endogenous staining. Scale bars: 20µm / 2µm in magnification. C: Co-immunoprecipitation experiment made in HEK293 cells expressing either GFP-KIAA1217 /p140Cap-Flag, GFP-KIAA1217/mito-Flag or GFP/mito-Flag. Cell extracts were immunoprecipitated with GFP or Flag traps. The input (I), the unbound fraction (FT) as well as the bound fraction (IP) were separated on SDS-PAGE and revealed with either rabbit polyclonal anti-GFP or rabbit polyclonal anti-Flag antibodies. Note that GFP-KIAA1217 immunoprecipitates p140Cap but very poorly mito-Flag. Similarly, p140Cap-Flag immunoprecipitates GFP-KIAA1217 but mito-Flag does not.

### Ciliogenesis defects triggered by either KIAA1217 or p140cap depletion are rescued by Src inhibition

Both KIAA1217 and p140Cap are located at the centrosome and regulate ciliogenesis. Since p140Cap is known as a Src inhibitor (Salemme et al., 2021b) and Src activity fine-tunes ciliogenesis^17,18^, we tested the hypothesis that both KIAA1217 and p140Cap regulate cilia formation via Src inhibition. We examined the potential rescue effect of two drugs on cilia formation in cells depleted for either KIAA1217 or p140Cap. Src family kinases are key regulators of morphology and epithelial integrity through regulation of actin dynamics and cellular adhesions. Cytochalasin D (CD) is known to allow BB migration in spread cells (Pitaval et al., 2010) and to enhance the centrosomal branched actin network, increasing the transport of preciliary vesicles to the mother centriole and promoting ciliation and cilia lengthening (Wu et al., 2018). Therefore, we first decided to analyze cilia formation in the presence of 0.5μM of CD in various siRNA conditions (control, KIAA1217 or p140Cap). As another control for ciliogenesis, we also depleted the protein MNR, which to our knowledge prevents ciliogenesis independently of actin (Chevrier et al., 2016; Kumar et al., 2021; Le Borgne et al., 2022). As expected, in control and MNR siRNA conditions, the presence of CD did not change the number of cilia but increased ciliary length in control condition (>6μm). CD treatment on cells depleted in either p140Cap or KIAA1217, increased significantly the number of cells bearing a cilium (Fig.9A-B) (for p140Cap, 80% ciliation with CD versus 20% ciliation without CD; for KIAA1217, 80% ciliation with CD versus 50% without CD). Cilia also appeared longer, both in p140Cap-and KIAA1217 depleted cells (Fig. 9C). Altogether, these results suggest that defective cilia formation after p140Cap or KIAA1217 depletion is respectively fully or partially rescued by CD treatment that promotes centrosomal branched actin network.

**Fig. 9:**
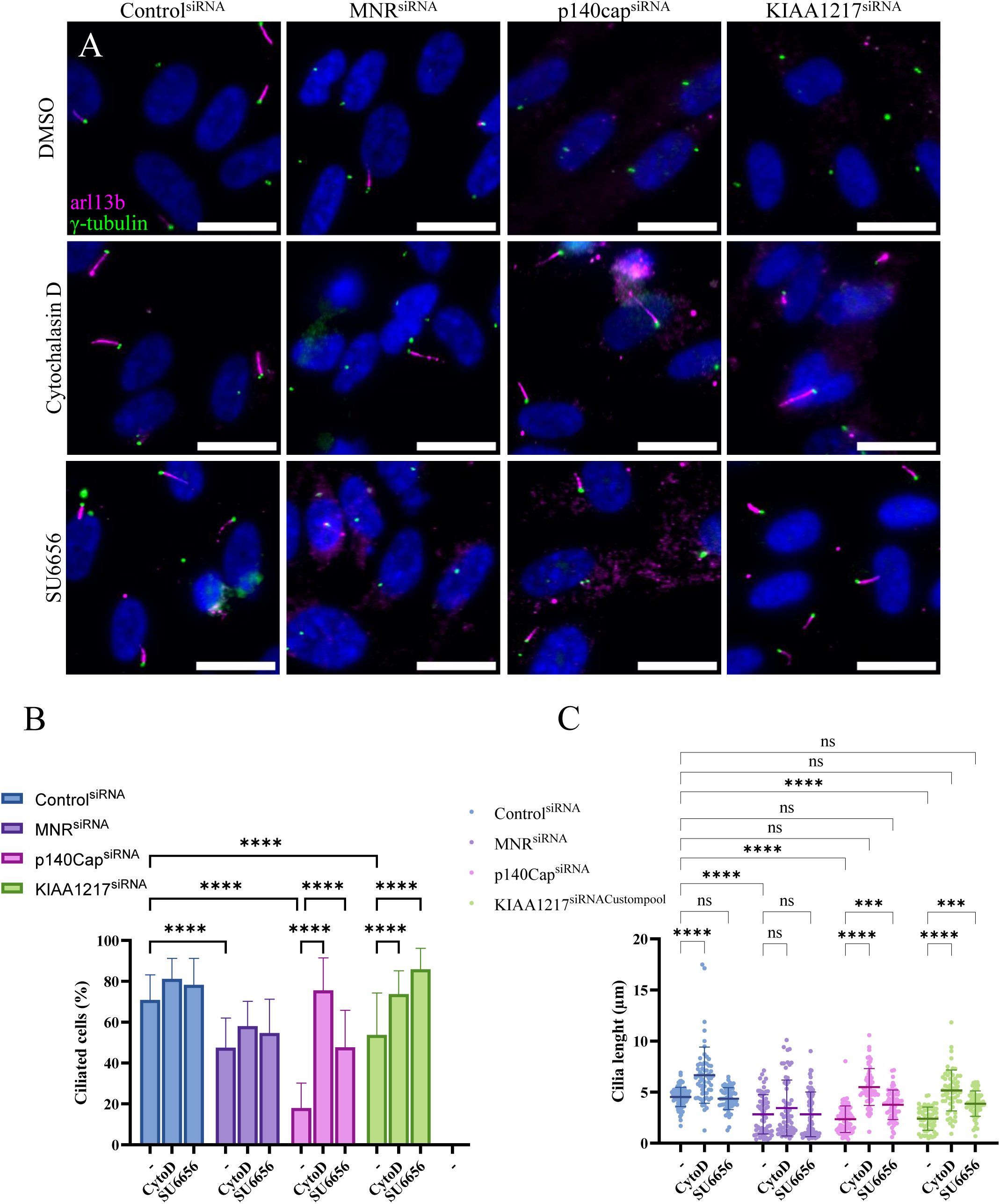
Actin or SRC inhibitors (CytochalasinD or SU6656) rescue ciliogenesis defect in p140Cap and KIAA1217 depleted cells. A: RPE-1 cells were transfected with either control^siRNA^, MNR^siRNA^, p140Cap^siRNA^ or KIAA1217^siRNACustompool^ in control condition (DMSO, upper lane), after cytochalasinD (middle lane) or SU6656 treatments (bottom lane). Representative images of the different depleted cells stained with both anti-arl13b and γ-tubulin antibodies 24 h after serum starvation and drug supplementation. Scale bars: 20µm. B: Quantification analysis of the percentage of ciliated cells after 24h of control treatment (DMSO), CytochalasinD (CytoD), or SU6656 (SU6656) treatments upon serum starvation in control-, MNR-, p140Cap-, KIAA1217 - depleted cells. The bar graph indicates the average from three independent experiments. Total cell number analyzed for each condition>539. All data are presented as average ± SD. ***p<0.001 / ****p <0.0001 (Two-way ANOVA followed by Tukey multiple comparisons test) C: Scatter plot representing the ciliary length (µm)after CD or SU treatment on control^siRNA^, MNR^siRNA^, p140Cap^siRNA^ and KIAA1217^siRNACustompool^. Cilia number analyzed in each condition >60 in two independent replicates. All data are presented as average ± SD. ***p<0.001 / ****p <0.0001 (Two-way ANOVA followed by Tukey multiple comparisons test)

Then, we decided to use the ATP-competitive inhibitor of Src, SU6656 (SU), which is known to inactivate different members of the Src family. As for CD experiment, the rescue of cilia formation was analyzed in cells inactivated in KIAA1217 or p140Cap in the presence of SU (1µM). Cilia formation either in control RPE-1 cells or in MNR siRNA-depleted cells was not modified by SU addition in the culture medium (Fig. 9A-C). In contrast, ciliation reached 48% in cells depleted for p140Cap and treated with SU versus 18% in cells without SU. Similarly, ciliation increased to 86% in depleted cells for KIAA1217 with SU versus 54% without SU (Fig. 9B). Interestingly, cilia length was not modified in control^siRNA^ or MNR^siRNA^ as opposed to CD treatment. However, both CD and SU treatments rescued ciliary length in both p140Cap and KIAA1217 depleted cells (Fig. 9C). This strengthens the hypothesis that p140Cap and KIAA1217 antagonize Src activity to facilitate ciliogenesis. Indeed, we propose that p140Cap and KIA1217 finely tune Src family activity at the BB to maintain the optimal actin dynamics.

## DISCUSSION

In this study using BioID/MS technology, we identified 24 CYLD proximity partners. As expected, centrosomal proteins and known CYLD partners were found such as SPATA 2, SPATA2L (Wagner et al., 2016; Schlicher et al., 2016, 2017), TRAF2, TRAF3 (Brummelkamp et al., 2003; Keats et al., 2007; Kovalenko et al., 2003; Trompouki et al., 2003) and CEP192 (Gomez-Ferreria et al., 2012). Despite the low number of identified CYLD-proximity partners, GO term analysis indicated that they are involved in ubiquitination, microtubule and cytoskeleton regulation in keeping with documented CYLD functions(Yang and Zhou, 2016). Among the candidates, KIAA1217 centrosomal protein (Gupta et al., 2015) was selected since its precise physiological function is still poorly elucidated, despite genetic data showing its involvement in axial skeletal malformation (Semba et al., 2006; Dai et al., 2024; Karasugi et al., 2009; Al Dhaheri et al., 2020), epithelial-mesenchymal transition (EMT) (Wang et al., 2021) and as a determinant of the a network of PSD proteins. Several arguments suggest that KIAA1217 and CYLD are involved in similar pathways. First, KIAA1217 and CYLD interact together and has synaptosomal SHANK3 as a common interactor (Jin et al., 2019; Morellato et al., 2025) Interestingly, *SKT*-KO (Morellato et al., 2025) and *CYLD-*KO mice (Colombo et al., 2021) show similar cognitive dysfunction, increased repetitive behavior and impaired excitability. Concerning the ciliogenesis, we and others have previously shown that CYLD regulates the ciliogenesis process (Eguether et al., 2014; Yang et al., 2014). Interestingly, CYLD reorganizes the actin network through the spatial activation of Src (Azimifar et al., 2012b). In agreement, we show that Src activation resulting from either KIAA1217 or p140Cap depletion prevents ciliogenesis by actin remodeling. Nevertheless, the molecular links between CYLD and KIAA1217 in ciliogenesis and synaptic plasticity remain to be elucidated in further studies.

### KIAA1217 localizes to the centrosome, microtubule +TIPS and FA

We demonstrate here, that the KIAA1217 protein localizes at the centrosome and microtubule + Tips at both endogenous and exogenous expression levels. Our results confirm the presence of eight SxIP/SxLP motifs in the KIAA1217 primary sequence, as previously reported (Jiang et al., 2012); (ii) Immunolabeling experiments revealed that KIAA1217 co-localizes with EB1 at endogenous and exogenous expression level; (iii) Live imaging experiments on GFP-KIAA1217 expressing cells exhibit comet-like structures emerging from the centrosome that are co-labelled with EB3-mcherry. (iv) disruption of the four C-terminus SxIP/SxLP motifs invalidates its EB1 binding ability(Jiang et al., 2012) and prevent the visualization of GFP-KIAA1217 comet-like structures by live imaging, even though EB3-mcherry still proved their presence. (v) Finally, GFP-KIAA1217 immunoprecipitates EB1-/SKT mcherry.

The presence of KIAA1217/SKT at both microtubule –ends and +Tips, as well as enrichments at FA is reminiscent of the Clasp family members subcellular distribution (Bouchet et al., 2016; Grimaldi et al., 2014; Mimori-Kiyosue et al., 2005; Stehbens et al., 2014). Indeed, Clasp proteins localize to the Golgi apparatus (Akhmanova et al., 2001), known as a microtubule nucleation center, and to the microtubule +Tips through a direct binding to the EB1 protein family (Mimori-Kiyosue et al., 2005), and to FAs (Bouchet et al., 2016; Stehbens et al., 2014). It is worth noting that the interaction of the Clasp proteins to the microtubule +Tips modulates microtubule dynamics (Aher et al., 2018; Lawrence et al., 2018; Mimori-Kiyosue et al., 2005; Patel et al., 2012; Straube and Merdes, 2007). Besides, clustering of CLASPs and microtubules around FAs temporally facilitates their disassembly by FA-associated extracellular matrix (ECM) degradation (Stehbens et al., 2014). Together, these data lead us to propose that KIAA1217 may modulate microtubule dynamics through its EB1 binding, FA turnover and extracellular matrix (ECM) degradation. To support a KIAA1217 function on FA turnover, the recent publication of Wang and collaborators (Wang et al., 2021) show that (i) KIAA1217 is overexpressed in hepatocellular carcinoma of bad prognostic caused by metastases; (ii) KIAA1217 overexpression promotes cell migration and invasion; and (iii) KIAA1217 overexpression increases the expression of the metalloproteases MMP2 and MMP9 (Wang et al., 2021); iv) KIAA1217 is involved in the EMT (Wang et al., 2021).

### KIAA1217 and p140Cap regulate ciliogenesis

This study mainly focused on the function of KIAA1217 in ciliogenesis. Previously, in a high throughput screen, KIAA1217 has been identified as a cilia positive regulator (Gupta et al., 2015). Here we show that siRNA-depletion of KIAA1217 in RPE-1 cells decreased cilia formation by one-third. In addition, about 70 % of the ciliated cells display very short cilia (less than 3μm). Interestingly, we also show that the siRNA-depletion of p140Cap, the KIAA1217 paralogue, drastically prevents cilia formation. Surprisingly, as KIAA1217 and p140Cap (Jaworski et al., 2009) can bind microtubules through EB1, we showed that modulating the actin network with CD treatment significantly increased cilia frequency and length after depletion of either p140Cap or KIAA1217, strongly suggesting that the two proteins regulate actin polymerization. Indeed, both microtubules and actin filaments are major actors in this process (Jewett et al., 2021b; Kim et al., 2010, 2015; Pitaval et al., 2017a) and their dynamics are tightly interconnected (Pitaval et al., 2017b). Furthermore, p140Cap is known to regulate Src signaling (Salemme et al., 2021b) by preventing Src activation through its binding to C-terminal Src kinase (Csk). In addition, an increased Src activity induced by MIM depletion prevents cilia formation (Bershteyn et al., 2010). Interestingly, MIM knockdown was shown to hyperphosphorylate cortactin (CTTN), which is known to promote actin polymerization. In our case, the treatment with the Src family inhibitor, SU6656, rescued the effects of both p140Cap and KIAA1217 depletion. Altogether, these results suggest that both KIAA1217 and p140Cap regulate ciliogenesis through modulation of Src family signaling and the actin network. How precisely KIAA1217 and p140Cap act on the actin network is not yet elucidated and will be analyzed in future studies. It is also interesting to point that KIAA1217 localizes at the FA. Of note, several major FA components including FAK, Paxillin, Vinculin and Talin proteins are associated with the BB in multiciliated cells to form a complex called ciliary adhesion that interacts with the actin cytoskeleton (Antoniades et al., 2014). In multiciliated cells, these ciliary adhesion complexes are assembled at the distal end of the basal foot, known to organize the polarized organization of apical microtubule lattice without affecting planar cell polarity (Antoniades et al., 2014). These ciliary adhesions connect both the microtubule and the actin network (Antoniades et al., 2014; Chatzifrangkeskou and Skourides, 2022). In our study, U-ExM microscopy on cells expressing prGFP-KIAA1217 displays two main localizations: (i) one at the subdistal end emerging from the mother centriolar wall. This localization is similar to numerous subdistal-appendages proteins (SDA) such as ninein, CEP170, Kif2a and p150glued/dynactin complex (Mazo et al., 2016; Chong et al., 2020) and in agreement with the localization of ciliary adhesion complex proteins at SDA (Chatzifrangkeskou and Skourides, 2022); (ii) In the second one, GFP-KIAA1217 forms a cap at the proximal end of both centrioles, reminiscent of PCM (pericentriolar material) proteins (Schweizer et al., 2021; Laporte et al., 2024) Altogether, these results suggest that KIAA1217 may function as a protein linking and regulating both the microtubule and the actin networks, reflecting the central role of the centrosome in organizing both microtubule and actin nucleation (Farina et al., 2019, 2016; Inoue et al., 2019; Obino et al., 2016).

### Involvement of KIAA1217 in spine skeletal malformation

Genetic studies performed in mouse report the involvement of KIAA1217 in intervertebral disc (IVD) malformations leading to tail curvature and in humans, mutations in KIAA1217 are associated either with lumbar disc herniation or vertebral malformations (Semba et al., 2006; Dai et al., 2024; Karasugi et al., 2009; Kelempisioti et al., 2011). Intervertebral discs (IVD) allow the spine to be flexible and to absorb shocks thus preventing vertebrae from grinding together. IVD comprises a gel-like nucleus pulposus (NP) in the center, concentric layers of collagen in the periphery, annulus fibrosus (AF) and a cartilaginous endplates (EP) that separate the IVD from the vertebrae. Interestingly, both cilia and ECM homeostasis appear important to preserve IVD properties. IVD degeneration is associated with a reduction in cilia length (Li et al., 2020). Postnatal loss of cilia in mice induces disc degeneration, through cells apoptosis (Li et al., 2020). IVD homeostasis also relies on FA proteins such as Kindlin-2 and integrin. The FA protein Kindlin-2 is highly abundant in nucleus pulposus (NP) and appears to regulate IVD homeostasis by activating the Nlrp3 inflammasome^69^. Intriguingly, a recent report suggests links between the centrosome and the Nlrp3 inflammasome through Spata2, CYLD and Nek7 protein (Chen et al., 2022). Integrin signaling via their ability to sense the environment, regulates cell death and ECM anabolism and catabolism (Chen et al., 2023). Interestingly, KIAA1217 is highly expressed in NP cells and a part of the AF (Abe et al., 2012; Semba et al., 2006). In HepG2 cells overexpressing KIAA1217, expression of matrix metalloproteinases (MMP2 and MMP9) is increased (Wang et al., 2021).

We speculate that KIAA1217 might be a central player in IVD formation and homeostasis, either via its role on cilia formation and homeostasis or via its function at FA. It would act as a bridge between cilia and inflammasome induction or acts through its regulation of FA function or MMP secretion to remodel the ECM.

In conclusion, our findings identify KIAA1217 as a novel regulator of ciliogenesis acting through Src-dependent modulation of the actin cytoskeleton. By bridging centrosomal, ciliary, and focal adhesion signaling, KIAA1217 appears as a central coordinator of cytoskeletal dynamics and cellular communication. Primary cilia and dendritic spines are cellular compartments specialized to sense and transduce environmental cues and presynaptic signals, respectively, while exhibiting similarities in many pathways and molecular mechanisms (Nechipurenko et al., 2013). The essential role of KIAA1217/SKT in these compartments suggests that this protein can represent a common component able to regulate membrane domain architecture, cellular interactions, and structural and functional plasticity both in cilia and in dendritic spines.

In the context of hepatocarcinoma metastasis, our results offer new insights into the molecular mechanisms of EMT driven by overexpressed KIAA1217, through its interaction with microtubule plus-ends via EB1 and its localization at focal adhesions, which could ultimately lead to better therapeutic strategies.

Furthermore, our work also opens new perspectives for investigating skeletal disorders associated with KIAA1217 mutations, providing a framework for better understanding the associated pathologies that currently lack clear molecular characterization. More broadly, these findings introduce new molecular players at the interface between ciliogenesis and intervertebral disc homeostasis, paving the way for future studies on cilia-related skeletal pathologies.

## MATERIAL AND METHODS

### Plasmids and reagents

All plasmids have been constructed using Gibson assembly and sequenced. Phusion DNA polymerase was used for all PCR.

GFP-CYLD and CYLD-Flag constructs were previously described (Eguether et al., 2014). Myc-BirA*CYLD was cloned into pcDNA5FRT/TO. All primers used in this study are listed in Supplementary Table 2.

Hs*Kiaa1217* cDNA was obtained from P. Defilippi laboratory. p140Cap-Flag was purchased from GenScript (clone ID: OHu23707), vectors EB1-GFP, EB3-mcherry were a gift of C. Janke (Curie Institute, Orsay, pTRIP-CAG-EB1-GFP, pTRIP-CAG-EB1-mcherry).

To invalidate Ile and Pro (IP) or Ile and Leu (LP) residues of the SxIP synthetic genes, shown in Supplementary Table 2, were ordered introducing the coding sequence for Asp (NN) instead of IP or LP. The KIAA1217 domains were subcloned in pEGFPC1.

For KIAA1217^siRNAcustom4Resistant^ plasmid, the sequence “AACCATCGATTGCTTCTA” was replaced by “AAGCCGAGCATCGCGAGT” in the Xba1-Xma1 fragment of pEGFPC1-KIAA1217.

### Cell culture and transfection

Human embryonic kidney (HEK293), human bone osteosarcoma (U-2 OS) cells were grown in DMEM supplemented with 10% of fetal bovine serum (FBS) and penicillin-streptomycin (1,000 units/mL and 1,000 μg/mL), and human retinal pigment epithelium (RPE-1) in DMEM/F12 with 2 mM L-glutamine. All cells were cultured in the presence of 5% CO2. Ependymal cells have been prepared as previously performed (Eguether et al., 2014).

Commercial Human glioblastoma (U-251) cells lines were cultivated in DMEM (Sigma) with 4.5 g/L L-glucose, L-glutamine, and pyruvate, supplemented with 10% FBS (Sigma) and 50 μg/mL gentamicin (Sigma).

For BioID experiment, Flp-In T-REx HEK293 cells were cotransfected with pOG44 (Flp-recombinase expression vector) and either pcDNA5FRT/TO mycBirA*-CYLD or pcDNA5FRT/TO mycBirA*. Transfections were performed with Lipofectamine2000 according to the manufacturer’s instructions. After transfection, cells were selected with hygromycin (200 μg/mL) and blasticidin (15 μg/mL). HEK293-FlpIn mycBirA*-CYLD and HEK293 FlpIn mycBirA* cells were incubated for 24 h with 50 μM biotin (Sigma-Aldrich) in the presence or absence of 0.1 μg/ml Tetracycline.

To establish RPE-1 cell line expressing GFP-KIAA1217 and GFP-KIAA1217^siRNAcustom4Resistant^, cells at 70% confluence were transfected with Lipofectamine2000 and 6μg of purified DNA plasmid (100mm dish). Transfected cells were selected using G418 at a concentration of 600µg/ml, until clones were observed. Clones were then isolated and analyzed by IF and WB for the GFP-KIAA1217 expression. Stable RPE-1 cell lines expressing GFP-KIAA1217 were grown on medium supplemented with G418.

Transient transfection in RPE-1 or U-2 OS was performed using lipofectamine2000 with 0.25 µg of plasmid DNA (12mm coverslip) for IF experiment according to the manufacturer’s protocol.

For IP experiments, transient transfection in HEK293 (100mm dish) was performed using polyplus transfection reagent with 10µg (7µg of plasmid expressing GFP-KIAA1217 and 3µg of plasmid expressing either CYLD-Flag, GFP, EB1-GFP, p140Cap-Flag) according to the manufacture’s protocol.

To induce cilia formation, RPE-1 cells (transfected or not) were seeded on coverslips until 100% confluence and incubated in starved condition (0.1% FBS medium) for 12-48 h. After IF and microscope image acquisition, at least 100 cells were counted on 10 fields randomly chosen for the quantification of ciliated cells for each experimental condition.

### BioID experiment

HEK293 mycBirA*-CYLD or HEK293 FlpIn mycBirA* cells were incubated for 24h with 50 μM biotin (Sigma-Aldrich) in the presence or absence of 0.1 μg/ml Tetracycline. Protein extraction for BioID purification was then performed according to(Gupta et al., 2015). Briefly, cell pellets were resuspended in lysis buffer (50mM Tris-HCl pH 7.5, 150mM NaCl, 1mM EDTA, 1mM EGTA, 1% Triton X-100, 0.1% SDS, 1:500 complete anti-protease inhibitor cocktail), and incubated at 4°C for 1 hour. The viscous lysate was passed through various needles of different diameters (0.6, 0.8 and 1.2mm) (Neolus™ Needles) until the lysate appeared liquid. After centrifugation at 16,000xg for 30 minutes at 4°C, supernatant was transferred to a fresh 15ml conical tube. 80μl of packed, pre-equilibrated Streptavidin sepharose beads (GE) were added and the mixture incubated overnight at 4°C with end-over-end rotation. Beads were pelleted by centrifugation at 800xg for 2 minutes and transferred with 1ml lysis buffer to a fresh 1.5ml Eppendorf tube. Streptavidin beads were washed 10 times with lysis buffer, and finally washed once with 50 mM Tris (pH 7.4). After the last wash, the proteins were recovered from the beads using Laemmli sample buffer. After incubation at 95°C for 5min, proteins were allowed to enter the SDS acrylamide gel for 1 cm. Gels were stained using colloidal blue and the gel part containing the proteins was then cut (gel plug). Three independent replicates were done for myc BirA* and mycBirA* CYLD cells.

#### Sample preparation prior to mass spectrometric analysis-

Individual bands of short SDS-PAGE migrations were excised for each sample, prepared in triplicate, and subjected to in-gel enzymatic digestion. Briefly, bands were washed sequentially with acetonitrile 30% and 100 mM ammonium bicarbonate. Proteins were reduced with 10 mM dithiothreitol (DTT) for 30 minutes at 56°C, followed by cysteine carbamidomethylation using 55 mM iodoacetamide (IAA) for 30 minutes. After supernatant removal, the washing steps were repeated, and the gel bands were dried. Tryptic peptides were generated by addition of sequencing-grade modified trypsin (5 ng/µl, Promega) and overnight incubation at 37°C. Proteolytic peptides were extracted first with 50% acetonitrile and 0.1% formic acid, then with 100% acetonitrile. The extracted peptides were vacuum-dried and resuspended in 2% acetonitrile with 0.05% trifluoroacetic acid (TFA) for nanoLC-MS/MS analysis

#### Mass spectrometry analysis-

NanoLC-MS/MS analysis was conducted using a nanoElute liquid chromatography system coupled to a timsTOF Pro2 mass spectrometer (Bruker, Billerica, MA, USA). Briefly, proteolytic peptides were desalted and preconcentrated online with a trap column (Waters, NanoE MZ Sym 18, 180 µm x 20 µm) and further separated onto an Aurora analytical column (ION OPTIK, 25 cm x 75 µm, C18, 1.6 µm), using a 0–35% solvent B gradient in 100 minutes. Solvent A was 2% acetonitrile and 0.1% formic acid in water, while solvent B was of 99.9% acetonitrile with 0.1% formic acid. MS and MS/MS spectra were recorded over a m/z range of 100 to 1700, with a mobility scan range of 0.65 to 1.45 V_s/cm². MS/MS spectra acquisition was performed using the PASEF (parallel accumulation–serial fragmentation) ion mobility-based acquisition mode, with the number of PASEF MS/MS scans set to 10.

#### Mass spectrometric Data analysis

Label-free quantification (LFQ) in MS1 was conducted with the MaxQuant software (version 1.6.6.0) using the LFQ algorithm combined with 4D feature alignment for a more specific quantitative analysis, including alignment of collisional cross-section (CCS) features through the « match between runs » function. Default normalization settings were applied, and protein identifications were performed using the Andromeda search engine against the UniProt human sequence database (downloaded on 08/04/19) supplemented with streptavidin and BirA* sequences, with default parameters. For the database search, following modifications were included: carbamidomethylation of Cysteins as fixed modification and Oxidation of Methionine as variable modification. MaxQuant protein group outputs were processed with the Perseus software (version 1.5.0. 0). Briefly, after log2 transformation of values, proteins detected in at least two replicates per condition were retained. Missing values were then imputed based on a normal distribution, before differential statistical analysis via Welch’s t-test. Proteins with significant changes in abundances were selected based on a p-value < 0.05 and fold-change (Fc) thresholds of Fc>2 or Fc<0.5 corresponding respectively to Log2Fc >1 and Log2Fc<-1. The mass spectrometry proteomics data have been deposited to the ProteomeXchange Consortium (http://proteomecentral.proteomexchange.org) via the PRIDE partner repository(Perez-Riverol et al., 2022) with the dataset identifier PXD 067058 and 10.6019/PXD067058.

### Home-made Antibodies

Antibodies against human KIAA1217 (rabbit) were produced against two C-terminal recombinant polypeptides EVITT-DTP (AA1339-1632) and NKF-KPT (AA1544-1791). 6×His-tagged EVITT-DTP and GST-NKF-KPT were produced and purified in BL21(DE3)pLysS bacteria and used to immunize animals. The purification of the antibodies was performed by antigen-specific affinity technique using the same polypeptides GST-EVITT-DTP or 6xHIS-tagged NKF-KPT. The specificity of the antibodies was tested by IF and WB in RPE-1cells depleted in KIAA1217. See Table 1 for antibodies used in this study.

### RNA interference

All siRNA were purchased from Dharmacon (horizondiscovery) (see Supplementary Table 2) and done as before(Le Borgne et al., 2022).

### Indirect immunofluorescence and microscopy

Cells grown on coverslips until confluence 70% were fixed in ice-cold methanol or after permeabilization with 0.5% Triton-X-100 and stained as previously described (Le Borgne et al., 2022).

All Figures presented are two-dimensional projections of images collected at all relevant z-axes using the confocal.

### Live imaging

Time-lapse image series were acquired with a 60x/1.49 NA Plan Apo objective lens on a microscope (Nikon Eclipse Ti) equipped with a spinning disk confocal scan head (CSUX1; Yokogawa) with a thermostatic humid chamber in 5% CO2 at 37°C and a camera (Prime95B, photometrix). Images of cells transfected with GFP-KIAA1217 with or without EB3 mcherry were acquired every 5 seconds for 5 minutes.

### TIRF Acquisition

Cells were plated on 1.5H ibidi glass bottom dishes (ref #81218-200). The following day immunostaining was performed as described in section “Indirect immunofluorescence and microscopy”. Wide field and azimutal TIRF acquisition were obtained using a Nikon eclipse Ti2 microscope

### Confocal acquisition

Confocal acquisitions were made with an inverted Leica SP8 or SP8x (for U-ExM) equipped with a UV diode (line 405) and 3 laser diodes (lines 488, 552, and 635) for excitation and 2 PMT detectors using a 63 × 1.4 NA oil objective. U-ExM images were deconvolved using water as a “mounting medium”. 3D stacks were acquired with 0.13 μm z-intervals and an x, y pixel size of 43 nm.

### FACS analysis

Cell cycle profile of control, KIAA1217, p140Cap and MNT-depleted serum-starved RPE-1 cells were determined by analyzing total DNA content using DAPI. Briefly, cells were trypsinized, fixed with 70% Ethanol at 4°C for 6 min, washed 3 times and resuspend in PBS. Cells were then labelled with a final concentration of 5 µg/ml 4’,6-diamidino-2-phenylindole (DAPI) and 10.000 cells were acquired with a Cytoflex S (Beckman-Coulter) cytometer, driven by Cytexpert 2.5 software. DAPI was excited by a 405-nm laser, and fluorescence was collected through a 450/45-nm band pass filter.

Data analysis was made with Kaluza 2.1 software (Beckman-Coulter). A first gate was made on an SSC-Area / DAPI-Area dot plot to select DAPI-labelled cells and doublets were discarded using a gate on DAPI-Area / DAPI-Height dot plot. Cell cycle was then analyzed on a DAPI-A distribution histogram using Michael H. Fox algorithm to determine frequency of G1-, S-and G2-phases.

The different conditions were compared to control condition using a Pearson’s Chi-squared Test with Package stats version 4.4.1 of R.

### Immunoprecipitations (IP)

Immunoprecipitations were carried out on HEK293 and same protocol was applied. Cells were lyzed in Tris 50mM, NaCl 150mM, 1%NP40 and complete inhibitor cocktail during 30 minutes at 4°C. Clear lysates were obtained after centrifugation 15 min at 13,000 rpm at 4°C. GFP trap and Flag trap beads (proteintech) were washed 3 times before use. Cell lysate was incubated with the beads for 3 h at 4°C under gentle agitation. Beads were washed 10 times with lysis buffer and finally resuspended in Laemmli sample buffer. After incubation at 95°C for 5min, proteins were separated by SDS-PAGE and process for WB.

### Centrosome purification

Centrosomes were isolated from KE-37 cells as previously described (Gogendeau et al., 2015). Briefly, cells were pretreated for 1 h with nocodazole (2 ×10−7 M) and cytochalasin D (2 × 10−6 M). Cells were then washed in PBS and resuspended in 8% sucrose in 10× diluted PBS before cell lysis in lysis buffer (1 mM HEPES, 0.5 mM MgCl2, 0.5% NP40, 1 mM phenylmethanesulphonyl fluoride and antiproteases). After centrifugation at 4000 rpm, the supernatant was filtered through a nylon mesh, readjusted to 10 mM HEPES and treated with DNAse and benzonase. Concentrated centrosomes were overlaid on a discontinuous sucrose gradient (70%, 50% and 40%) in a SW32Ti tube and centrifuged at 25 000 rpm for 1 h 15 min. Fractions were then collected and analyzed for their purity.

### Ultrastructural expansion microscopy (U-ExM)

For analyzing KIAA1217 in human centrioles, U-ExM was performed as previously described (Gambarotto et al., 2019). Briefly, cells grown on 12mm coverslips were incubated in 2% AA + 1.4% FA diluted in PBS for 3 to 5 h at 37°C prior to gelation in monomer solution (19% sodium acrylate, 0.1% bis-acrylamide, and 10% acrylamide) supplemented with TEMED and APS (final concentration of 0.5%) for 30 min to 1 h at 37°C. Denaturation was performed for 1 h 30 min at 95°C. After 2×20 min of washes in ddH2, gels were measured with a millimeter paper. Measurements of lengths and diameters were scaled according to the expansion factor of each gel. Gels were placed in PBS for 15 min and incubated with primary antibodies in PBS-BSA 2% for 2h30 at 37°C.

### Drug treatments

RPE-1 cells were treated with either control^siRNA^, MNR^siRNA^, p140Cap^siRNA^ or KIAA1217^siRNA^. After 48h of depletion cells were confluent and induced for ciliation by lowering the serum to 0,1% in the culture medium. At that time, cells were treated with either cytochalasinD at 0.5µM (Kim et al., 2010) or SU6656 at 1µM (Bershteyn et al., 2010) for 20h before fixation.

---

**Fig. S1:**
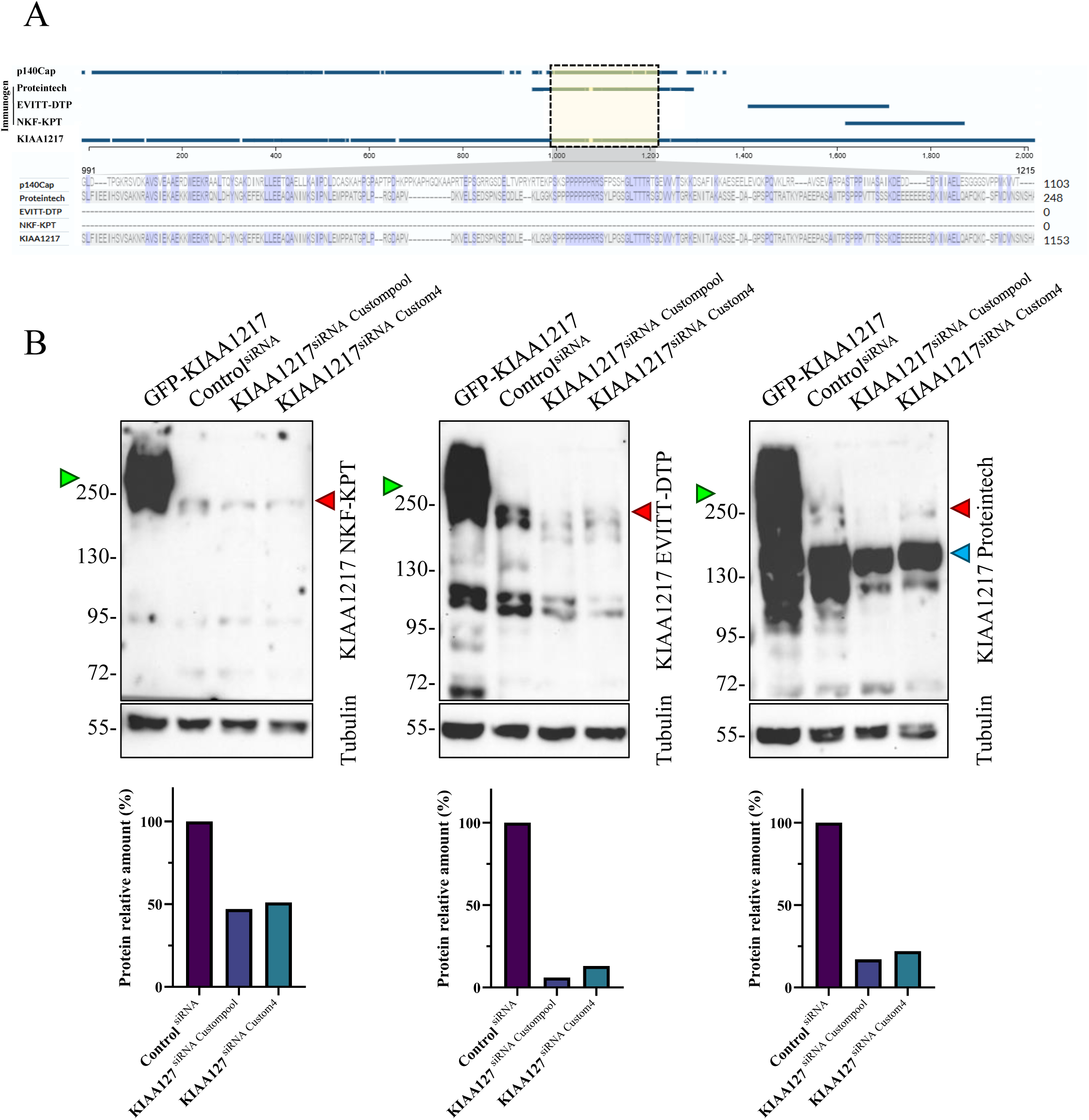
KIAA1217 antibodies validation. A: Graph showing an alignment of p140Cap and KIAA1217 with a blow-up of the AA991-1215 in the yellow square. The different immunogens (proteintech, EVITT-DTP and NKF-KPT) are positioned on this alignment. B: Three WB revealed by anti-KIAA1217 EVITT-DTP, anti-KIAA1217 NKF-KPT and the Proteintech antibodies. The different lysates loaded on the gels are indicated on the figure. The green arrowhead indicates the GFP-KIAA1217 band, the red arrowhead endogeneous KIAA1217 and the blue arrowhead the major band recognized by the KIAA1217 proteintech antibodies. Anti-Tubulin antibody is used as loading control. The endogeneous KIAA1217 band is diminished by KIAA1217^siRNA^. The graphs represent the luminescence quantification of the KIAA1217 depletion by siRNA. In each conditions, KIAA1217 intensities are divided by those of tubulin and all data are normalized base on each control siRNA.

**Fig. S2:**
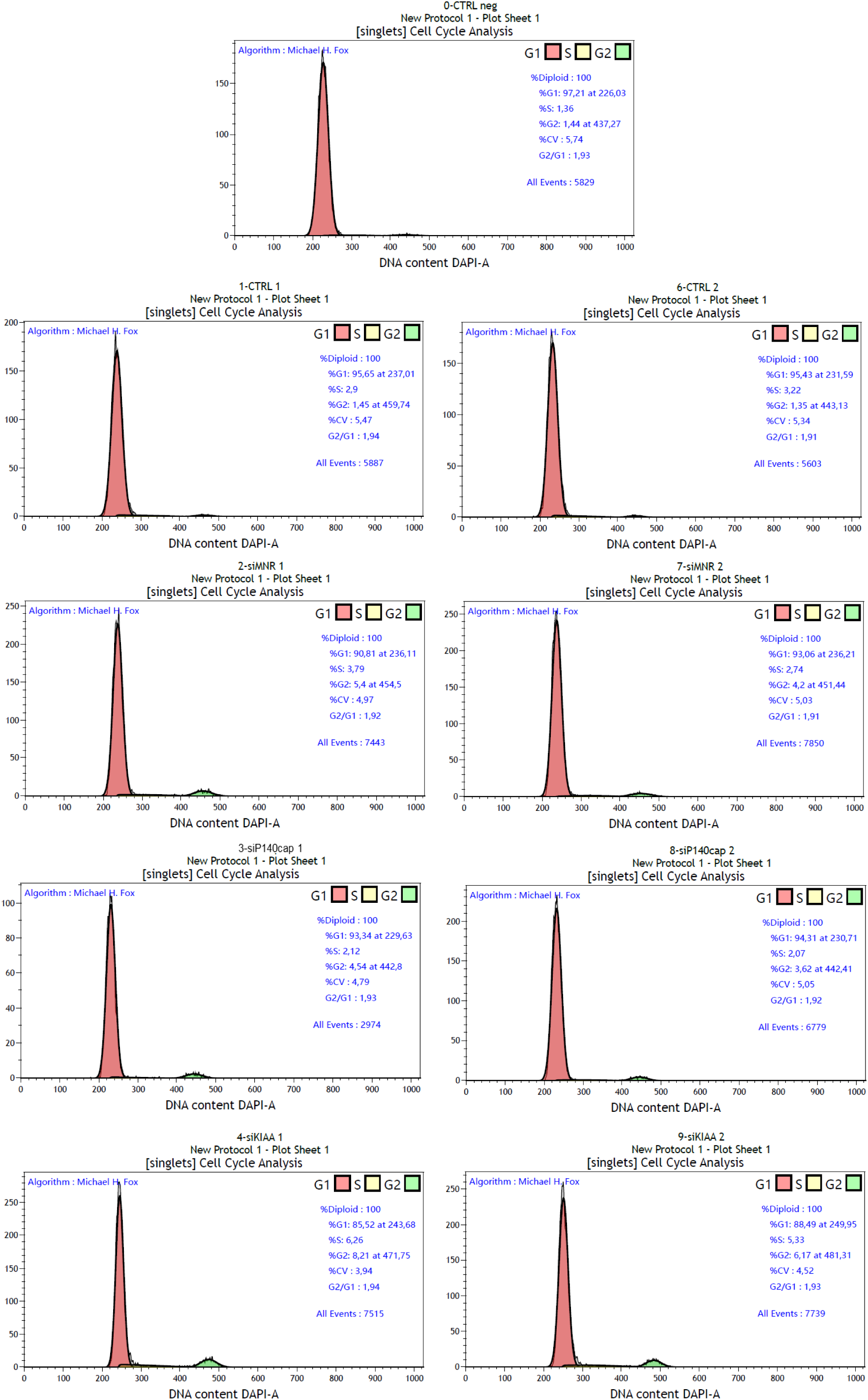
cell cycle analysis. Cell cycle analysis of RPE-1 cells after serum starvation and treatment with control^siRNA^, MNR^siRNA^, P140Cap^siRNA^and KIAA1217^siRNA^.

**Fig. S3:**
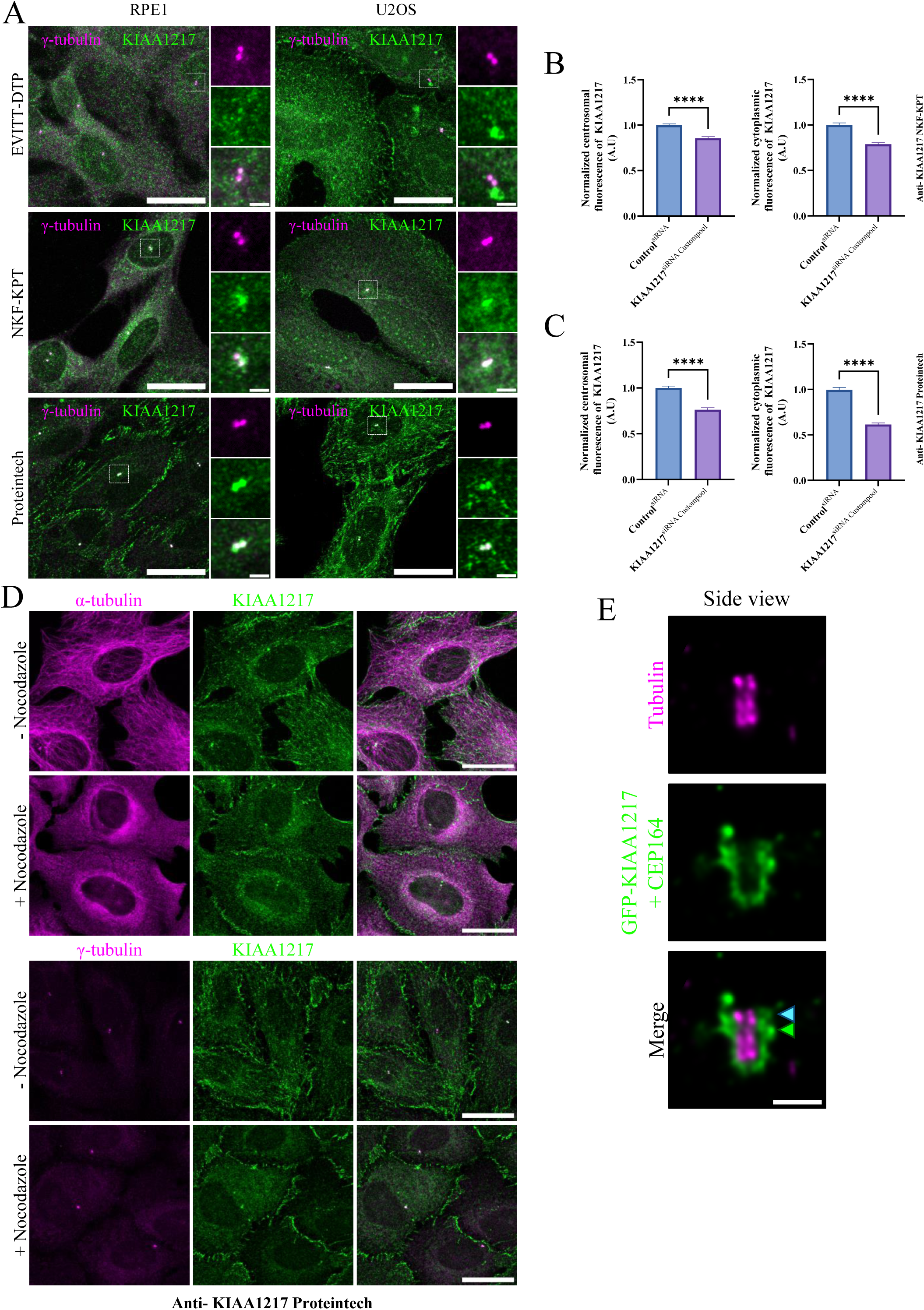
Validation of the three KIAA1217 antibodies by IF. A: Immunolabeling of either RPE-1 or U2-OS cells with the three different KIAA1217 antibodies. All antibodies recognize the centrosome. Scale bars: 20µm / 2µm in magnification. B-C: Graph showing the Quantification of the KIAA1217 fluorescence at the centrosome and the cytoplasm in control^siRNA^ (dark grey) or KIAA1217^siRNACustompool^ (light grey) as revealed by anti proteintech (B) or the KIAA1217 NKF-KPT (C) antibodies. The bars show the average intensity of fluorescence (n > 40cells, 2 independent replicates). Error bars show SD. Statistical significance was assessed by one-way ANOVA followed by Tukey’s post hoc test ****p < 0.0001. D: Immunolabeling of U-2 OS cells with anti-KIAA1217 (Proteintech) and α-tubulin or γ-tubulin without or with Nocodazole. Scale bars: 20µm. E: U2-OS co-stained for CEP164 and GFP-KIAA1217 (green) and tubulin (magenta) centrosome observed by U-ExM. The image has not been deconvoluted. The light blue arrowhead indicates the CEP164 staining and the green one the GFP staining. Scale bars: 250nm

**Fig. S4:**
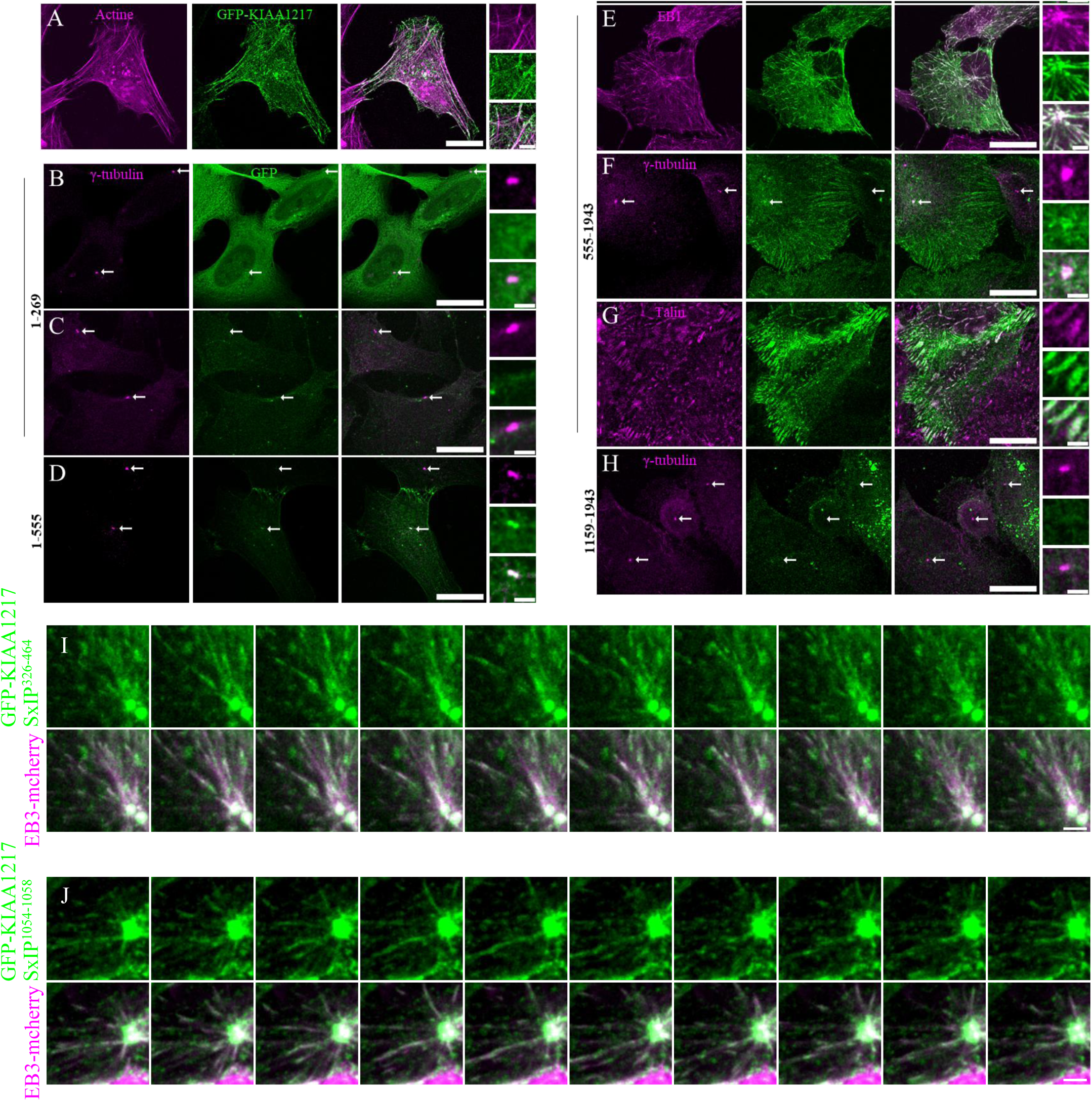
expression of constructs producing truncated KIAA1217 and mutated SxIP domains. A: U2-OS cells expressing full-length GFP-KIAA1217 after glutaraldehyde fixation stained with phalloidin. Some actin filaments are decorated by GFP-KIAA1217. Scale bars: 20µm and 2µm for the magnifications. B-H: Localization of the different constructs of KIAA1217 tagged with GFP AA1-269 (B,C); AA1-555 (D); AA555-1943 (E-G) and AA1159-1943 (H) after Methanol fixation without permeabilization (B,E) or after triton X-100 extraction(C,D,F,G,H). Cells were stained for GFP (B-H) and γ-tubulin (B-D,F,H), EB1(E), and Talin (G). Scale bars: 20µm / 2µm in magnification. I-J: Localization of GFP-KIAA1217 regarding the microtubule +Tips protein (EB1 or EB3) Series of 10 images were acquired every 5 seconds of RPE-1 cells expressing GFP-KIAA1217 with mutated SxIP^326-464^ (I) SxIP^1054-^1058 (J) with EB3 mcherry. Comet-like structures revealed by GFP auto-fluorescence emerging from the centrosome is followed through the movie and colocalized with EB3mcherry (auto-fluorescence). Scale bars: 2µm.

**Fig. S5:**
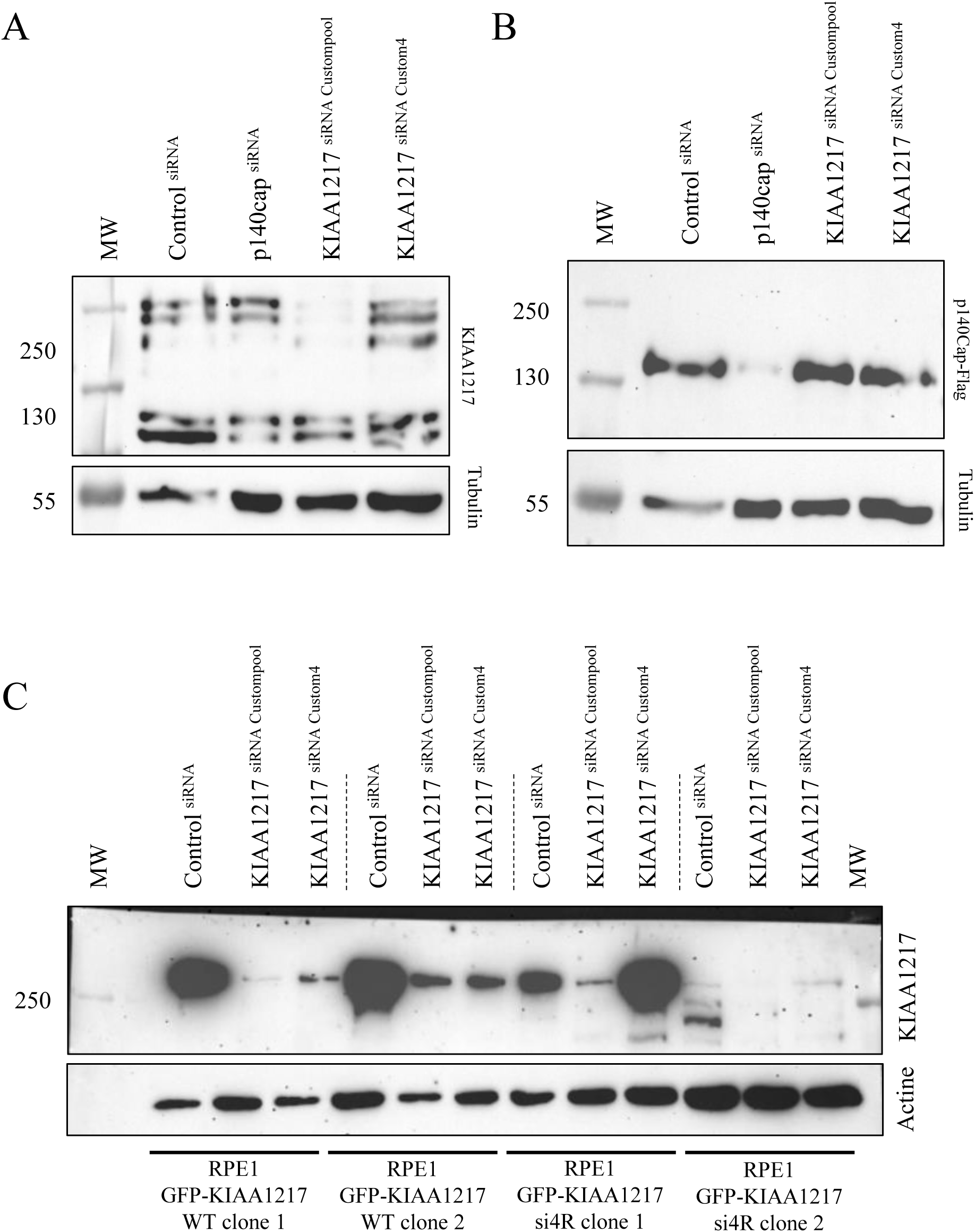
Efficiency of p140Cap^siRNA^ and KIAA1217^siRNA^. A: WB revealed by anti-KIAA1217antibodies (Proteintech) of RPE-1 cells depleted with: control^siRNA^, p140Cap^siRNA^, KIAA1217^SiRNA^C^ustom^ ^pool^ or KIAA1217 ^siRNA^ ^Custom4^ and anti-Tubulin antibodies as loading control. Note that the expression of KIAA1217 is not affected by the depletion of p140Cap, whereas KIAA1217 ^SiRNACustom^ ^pool^ or KIAA1217 ^siRNA^ ^Custom4^ decrease its expression. B: WB revealed by anti-Flag antibodies of RPE-1 cells expressing p140Cap-Flag transiently and depleted with: control^siRNA^, p140Cap^siRNA^, control^siRNA^, KIAA1217^SiRNACustom^ ^pool^ and KIAA1217^siRNA^ ^Custom4^ and anti-Tubulin antibodies as loading control (MW: molecular weight marker). Note that the expression of p140Cap-Flag is only decreased by p140Cap^siRNA^ and not by KIAA1217^SiRNACustompool^ and _KIAA1217_siRNACustom4. C: WB of stable RPE-1 cell lines overexpressing GFP-KIAA1217 WT (clone1 and 2) and GFP-KIAA1217^siRNA4^ ^resistant^ (clone 1 and 2) and treated with either control^siRNA^, KIAA1217^siRNACustom^ ^pool^ or KIAA1217^siRNACustom4^. The blot is revealed with anti-KIAA1217 (Proteintech) and anti-Tubulin antibodies as loading control.

***Supplementary movie1***: Time-lapse of RPE-1 cells expressing GFP-KIAA1217 (green) and EB3-mcherry (magenta) corresponding to Fig. 7C.

***Supplementary movie2***: Time lapse of RPE-1 cells expressing GFP-KIAA1217 SxIP^1799-1855^ (green) and EB3-mcherry (magenta) corresponding to Fig. 7D

***Supplementary movie3***: Time lapse of RPE-1 cells expressing GFP-KIAA1217 SxIP^326-464^ (green) and EB3-mcherry (magenta) corresponding to Supplementary Fig. S4

***Supplementary movie4***: Time lapse of RPE-1 cells expressing GFP-KIAA1217 SxIP^1054-1058^ (green) and EB3-mcherry (magenta) corresponding to Supplementary Fig. S4

## Abbreviations

BB: basal body
FA: focal adhesion
ECM: extracellular matrix
EMT: epithelio-mesenchymal transition
BioID: In vivo biotinylation identification
MS: mass spectrometry
SKT: sickle tail
GFP: green fluorescent protein
aPKC: atypical protein kinase C
MIM: Missing-in-Metastasis
PKD: polycystic kidney diseases
ARPKD: autosomal recessive polycystic kidney diseases
ADPKD: autosomal dominant polycystic kidney diseases
ILK: integrin-linked-kinase
EGF: epidermal growth factor
SU: SU6656
CD: CytochalasinD
IF: immunofluorescence
WB: Western blot
U-ExM: ultrastructural expansion microscopy
ATP: adenosin triphosphate
KO: knock-out
IVD: intervertebral disc
NP: nucleus pulposus
AF: annulus fibrosus
EP: endplate
MMP: matrix metalloproteinase

## Acknowledgments

We would like to thank Carsten Janke for his generous gift of anti-poly-E antibodies and for EB1 and EB3 plasmids. We also would to thank Cyndy Mathon for protein purification which allowed both the antigen preparation and affinity purification of the antibodies and Karima Palmier for her help in plasmid cloning. We thank M. Laporte, T. Eguether, C. Vesque, P. Guichard, for critical reading of the manuscript. The present work has benefited from Imagerie-Gif core facility supported by Agence Nationale de la Recherche (ANR-10-INBS-04/FranceBioImaging; ANR-11-IDEX-0003-02/ Saclay Plant Sciences). We acknowledge the Proteomic-Gif (SICaPS) platform (supported by IBiSA, Ile de France Region, PlanCancer, CNRS and Paris-Saclay University). This work was supported by the CNRS (UMR9198), LG was funded by a PhD fellowships from Université Paris-Saclay (https://www.paris-saclay.fr) and FRM (FDT202304016625).

## Funding

This work is supported by Ligue Nationale contre le Cancer (M35345) to AMT and Agence Nationale de la Recherche (ANR-22-CE13-0014-03) to AMT.

## Author contributions

L.G. performed most of the experiments, analyzed the data, and prepared the figures. L.R. performed co-expression experiments followed by immuno-precipitations. H.G. and S.E-M performed TIRF microscopy. J. B. designed and cloned the BirA*-CYLD plasmid, established the cell line and performed BioID preliminary experiments. P.D. send us the Hs*Kiaa1217* cDNA. L.S. and V.R. performed the MS and analyzed the results. K.B. and A-M T. performed the BioID experiment and analyzed the results. KB performed some clonings. A-M T. designed the experiments, supervised the project, and wrote the manuscript with L.G. All authors review and edit the manuscript.

The authors declare that they have no competing interests.

## Notes

### Competing Interest Statement

The authors have declared no competing interest.

### Summary of Updates

The title has been changed to reflect our findings.

