## Supplementary Table 1 for "KIAA1217/SKT and its paralog p140Cap are centrosomal proteins that regulate ciliogenesis through Src family signaling pathway"

### Supplementary Table1

#### Primary antibodies

| Primary antibodies | | reference | species | IF dilution | WB dilution | expansion microscopy |
|---|---|---|---|---|---|---|
| KIAA1217 EVITT-DTP | Home made | #1 | rabbit | too weak | 1/1000 | |
| KIAA1217 NKF-KPT | home made | #2 | rabbit | 1/500 | 1/500? | |
| KIAA1217 | Proteintech | 24880-1-AP | rabbit | 1/500 | 1/1000? | |
| p140Cap | Saint John's laboratory | STJ94856 | rabbit | 1/500 | | |
| EB1 | BD transduction laboratory | 610535 | mouse | 1/1000 | 1/1000 | |
| tubulin clone B5.1.2 | Sigma | T5168 | mouse | 1/1000 | 1/1000 | |
| talin | Proteintech | 14168-1-AP | rabbit | 1/250 | 1/1100 | |
| paxillin | BD Transduction | 610057 | mouse | 1/500 | | |
| Arl13 | Clinisciences, clone N295B/66 | Ab02312-2.0 | mouse | 1/400 | | |
| anti-γ-tubulin clone GTU-88 | Sigma | T6557 | mouse | 1/1000 | | |
| anti-GFP | Torrey Pines | TP-401 | rabbit | | | 1/400 |
| anti-GFP | Institut curie | | rabbit | 1/1000 | 1/1000 | |
| anti-Flag | Proteintech | 20543-1-AP | rabbit | 1/1000 | 1/1000 | |
| anti-Flag | Sigma -M2 | F4049 | mouse | 1/1000 | 1/1000 | |
| Mouse monoclonal alpha-tubulin | ABCD antibodies | AA345 | mouse | | | 1/250 |
| Mouse monoclonal beta-tubulin | ABCD antibodies | AA344 | mouse | | | 1/250 |
| GT335 | Gigft C. Janke | | mouse | 1/10000 | | |
| poly-E | Gigft C. Janke | | rabbit | 1/1000 | | |

#### Secondary antibodies

| Secondary antibodies | | reference | species | IF dilution | WB dilution | expansion microscopy |
|---|---|---|---|---|---|---|
| Alexa fluor 488 F(ab')2 frgt anti rabbit IgG | Invitrogen | A11070 | goat | 1/1000 | | 1/250 |
| Alexa fluor 647 anti rabbit IgG | Invitrogen | A21244 | goat | 1/1000 | | |
| Alexa fluor 488 anti mouse IgG | Invitrogen | A11029 | goat | 1/1000 | | |
| Alexa fluor Plus 647 anti mouse IgG | Invitrogen | A32728 | goat | 1/1000 | | |
| Alexa fluor 568 anti mouse IgG | Invitrogen | A11031 | goat | 1/100 | | 1/250 |
| anti-mouse immunoglobulins peroxidase | Sigma | A0412 | goat | | 1/10000 | |
| anti-rabbit immunoglobulins peroxidase | Sigma | A0545 | goat | | 1/10000 | |

### Reagent and ressources

#### Expansion microscopy

| | | |
|---|---|---|
| Acrylamide (AA, 40%) | SIGMA | A4058 |
| Nuclease-free water | Ambion-ThermoFisher | AM9937 |
| Sodium Acrylate (97-99%) (SA) | SIGMA | 408220 |
| Ammonium persulfate (APS) | ThermoFisher | 17874 |
| Tetramethylethylendiamine (TEMED) | ThermoFisher | 17919 |
| N,N'-methylenbisacrylamide (BIS) 2% | SIGMA | M1533 |
| PolyDLysine | Gibco | A3890401 |

#### cell cultures reagent

| | | |
|---|---|---|
| DMEM glutamax | Fisher scientific | 11584486 |
| DMEM/F12 | Fisher scientific | 11540446 |
| RPMI 1640 | Fisher scientific | 11560586 |
| fetal calf serum | | |
| penicilline steptomycine | Fisher scientific | 11556461 |
| L-glutamine | Fisher scientific | 15430614 |
| trypsin (o,25%) | Fisher scientific | 11560626 |
| lipofectamine 2000 | Fisher scientific | 12313563 |
| lipofectamine RNAimax | Fisher scientific | 12343563 |
| jetprime | ozyme | POL101000046 |
| cytochalasinD | Sigma | C8273 |
| SU6656 | Sigma | 572635 |
| Complete anti-proteases | Roche | |

#### Software and algorithms

| | |
|---|---|
| graphPad Prism | GraphPad Software,LLC |
| PickCentrioleDim' | Laporte et al., 2024 |
| centrioleGraph | Laporte et al., 2024 |
