## Supplementary Table 2 for "KIAA1217/SKT and its paralog p140Cap are centrosomal proteins that regulate ciliogenesis through Src family signaling pathway"

| Supplementary Table2 | | | |
|---|---|---|---|
| **Constructs** | **primers used 5'-3'** | | |
| pEGFP C1 KIAA1217 FL | GFP :TCAGATCCGCTAGCGCTACCGGTCGCCACCATGGTGAGCAAGGGCGAGGAGCTGTTCACC and<br>CTCACATTTCTGGCTTTCATTTTCTTCCATgccccctccaccgccCTTGTACAGCTCGTCCATGCCGAGAGTGA<br>KIAA1217: ATGGACGAGCTGTACAAGggcggtggagggggcATGGAAGAAAATGAAAGCCAGAAATGTGAGCC and<br>TATGATCAGTTATCTAGATCCGGTGGATCTTAAGAGGTTTCTTTTGCTGTGGATGGGGTG | | |
| pEGFP C1 KIAA1217 FL siRNA custom4 resistant | GFP idem as FL and | | |
| pEGFP C1 KIAA1217 1-269 | GFP idem as FL and | | |
| pEGFP C1 KIAA1217 1-555 | GFP idem as FL | | |
| pEGFP C1 KIAA1217 1-1159 | GFP idem as FL | | |
| pEGFP C1 KIAA1217 555-1943 | GFP idem as FL | | |
| pEGFP C1 KIAA1217 1159-1943 | GFP idem as FL | | |
| **SxIP mutation in SxNN** | **Synthetic genes 5'-3'** | | |
| pEGFPC3-KIAA1217/SKT-SxIP$^{326-464}$ | GAGCCTTCCTGTGGTGAGGTCAACCAACCAGACGAAAGAAagatctCTGGGGGTTCTCTATCTCCAGTATGGAGAT<br>GAAACCAAGCAGCTCAGGATGCCGAATGAAATCACAAGTGCAGACACAATCCGTGCTCTCTTCGTAAGTGCCTTT<br>CCACAGCAGCTCACCATGAAAATGCTGGAATCGCCCAGTGTCGCCATTTACATCAAAGATGAAAGCAGAAATGTC<br>TATTATGAATTAAATGATGTAAGGAACATTCAAGACAGATCACTCCTCAAAGTGTACAACAAGGATCCTGCACAT<br>GCGTTTAATCACACACCAAAAACTATGAATGGAGACATGAGGATGCAGAGAGAACTTGTTTATGCAAGAGGAGA<br>TGGCCCTGGGGCCCCTCGCCCCGGATCTACTGCTCATCCACCCCATGCGATTCCAAATTCCCCACCGTCTACTCCA<br>GTGCCCCATTCCATGCCCCCCTCCCCGTCCAGAaaTaaTTATGGGGGCACCCGCTCCATGGTTGTTCCTGGCAATG<br>CCACCATCCCCAGGGACAGAATCTCCAGCaacaatGTCTCCAGACCCATCTCTCCAAGCCCAAGCGCCATTTTAGAA<br>AGAAGAGATGTCAAGCCTGATGAAGACATGAGTGGCAAAAACATTGCAATGTACAGAAATGAGGGTTTCTATGC<br>TGATCCTTACCTTTATCACGAGGGACGGATGAGCATAGCCTCATCCCATGGTGGACACCCACTGGATGTCCCCGA<br>CCACATCATTGCATATCACCGCACCGCCATCCGGTCAGCGAGTGCTTATTGTAACCCCTCAATGCAAGCGGAAAT<br>GCATATGGAACAATCACTGTACAGACAGAAATCAAGGAAATATCCGGATAGCCATaacaaTACACTGGGCTCCAA<br>AACACCCCCTGCCTCTCCTCACAGAGTCAGTGACCTGAGGATGATAGACATGCACGCTCACTATAATGCCCACGG<br>CCCCCCTCACACCATGCAGCCAGACCGGGCCTCTCCGAGCCGCCAGGCCTTTAAAAAGGAGCCAGGCACCTTGGT<br>GTATATAGAAAAGCCACGGAGCGCTGCAGGATTATCCAGCCTTGTAGACCTCGGCCCTCCTCTAATGGAGAAGC<br>AAGTTTTTGCCTACAGCACGGCGACAATACCCAAAGACAGAGAGACCAGAGAGAGGATGCAAGCCATGGAGAA<br>ACAGATTGCCAGTTTAACTGGCCTTGTTCAGTCTGCGCTTTTTAAAGGGCCCATTACAAGTTATAGCAAAGATGC<br>GTCTAGCGAGAAAATGATGAAAACCACAGCCAACAGGAACCACACAGATAGTGCAGGAACGCCCCATGTGTCTG<br>GTGGGAAGATGCTCAGTGCTCTGGAGTCCACGGTGCCTCCCAGCCAGCCTCCACCTGTGGGCACCTCAGCCATCC<br>ACATGAGCCTGCTTGAGATGAGGCGGAGCGTGGCGGAACTCAGGCTCCAGCTCCAGCAGATGCGGCAGCTCCA<br>GCTGCAGAACCAGGAGTTGCTGAGGGCAATGATGAAGAAGGCCGAGCTGGAAATCAGTGGCAAAGTGATGGA<br>AACAATGAAGAGACTGGAGGATCCCGTGCAGCGACAGCGCGTCCTAGTGGAGCAAGAGAGACAAAAATATCTT<br>CATGAGGAAGAGAAGATCGTCAAGAAGTTGTGCGAGTTGGAAGACTTTGTTGAAGACTTGAAGAAGGACTCCA<br>CGGCAGCCAGCCGATTGGTTACTCTGAAAGACGTGGAAGACGGGGCTTTCCTCCTGCGTCAAGTGGGAGAGGCT<br>GTAGCTACCCTGAAAGGAGAATTTCCAACCTTACAAAACAAGATGCGAGCCATCCTGCGCATAGAAGTGGAGGC<br>CGTGCGGTTTCTGAAGGAGGAGCCACACAAGCTGGACAGTCTCCTGAAGCGTGTGCGCAGCATGACAGACGTCC<br>TGACCATGCTGCGGAGACATGTCACTGATGGGCTCCTGAAAGGCACGGACGCAGCCCAAGCCGCACAGTACATG<br>GCTATGGAAAAGGCCACAGCCGCAGAAGTCCTGAAGAGTCAGGAGGAGGCAGCCCACACCTCCGGCCAGCCCTT<br>[illegible] | | |
| pEGFPC3-KIAA1217/SKT-SxIP$^{1054-1058}$ | TTTGTCCGGCATGATGGTTCGCCACGCGCAgagctcCCCTGTGGTCATCCAGCCCTCCCAGCACTCCGTGGCCCTGC<br>TGAACCCTGCTCAGAACTTGCCTCACGTGGCCAGCTCCCCAGCCGTCCCCCAGGAAGCAACCTCCACTCTGCAGAT<br>GTCGCAGGCTCCGCAGTCCCCACAGATACCCATGAATGGGTCTGCCATGCAGAGCTTGTTCATTGAAGAAATCCA<br>CAGTGTGAGTGCCAAGAACAGGGCAGTGTCTATCGAGAAAGCAGAAAAGAAATGGGAGGAAAAAAGGCAAAA<br>TCTGGATCACTATAATGGGAAAGAGTTTGAGAAGCTCCTAGAAGAAGCTCAGGCCAATATCATGAAGTCAATAC<br>CAAATCTGGAGATGCCGCCAGCCACAGGCCCACTGCCAAGGGGAGATGCCCCAGTGGACAAGGTGGAACTTTCA<br>GAAGATTCTCCAAATTCGGAACAGGACTTGGAAAAGCTGGGGGGAAAGTCGCCCCCTCCTCCTCCGCCACCTCCT<br>CGTCGAAGCTACaacaatGGATCGGGACTCACCACCACGAGGTCAGGCGATGTGGTCTACACCGGCAGAAAGGAG<br>AACATCACCGCTAAGGCAAGCAGTGAAGATGCTGGACCAAGCCCACAGACCAGAGCTACAAAATATCCAGCAGA<br>GGAGCCTGCTTCAGCCTGGACCCCATCCCCACCGCCTGTCACCACCTCCTCCTCAAAGGATGAGGAGGAAGAAGA<br>AGAAGAAGGAGACAAAATAATGGCAGAACTCCAGGCATTCCAGAAGTGTTCCTTTATGGATGTAAATTCAAACA<br>GTCATGCTGAGCCATcccgggCTGACAGTCACGTTAAAGACACTAGGTCGG | | |
| pEGFPC3-KIAA1217/SKT-SxIP$^{1799-1884}$ | CCCAAGAGGCCTCTCCCCGACCCTTGCTAGTTCCGGATGAaggtccCACTGCCCTAGAGCCCCCTACGTCGATACCT<br>TCAGCTTCACGTAAGGGCTCCAGCGGGGCCCCACAGACGAGCAGGATGCCTGTCCCCATGAGTGCCAAGAACAG<br>ACCCGGAACCCTGGACAAACCCGGCAAGCAGTCCAAACTGCAGGATCCCCGCCAATATCGTCAGGCTAATGGAA<br>GTGCTAAGAAATCTGGTGGGGACTTTAAGCCTACTTCCCCCTCCTTACCTGCTTCTAAGAaTaacGCCCTTTCTCCC<br>AGCTCTGGGAAAAGCAGTTCTaataaCTCTTCTAGTGGTGACAGCTCTAACaaCaaTAATCCACCTGCTACTAAACC<br>ATCGATTGCTTCTAACCCTCTCAGCCCCCAAACAGGACCACCTGCTCACTCTGCCTCCCTCAaCaaTTCTGTCTCTAA<br>TGGCTCTTTGAAGTTTCAGAGCCTCACTCATACAGGTAAAGGTCACCATCTTTCATTCTCACCGCAGAGTCAAAAT<br>GGCCGAGCACCCCCTCCTTTGTCATTTTCCTCCTCCCCTCCTTCTCCTGCCTCCTCCGTCTCACTGAATCAAGGTGCC<br>AAGGGCACCAGGACCATCCATACTCCCAGCCTCACCAGCTACAAGGCACAGAATGGAAGTTCAAGCAAAGCCAC<br>CCCATCCACAGCAAAAGAAACCTcttaagATCCACCGGATCTAGATAACTGATCATAATCAG | | |
| **siRNA** | | | |
| KIAA1217 siRNA custom | Horizondiscovery (Dharmacon) | siRNA KIAA1217/SKT 1 : 5'-CCAUGAAGAUGUACGGAAAA-3' | |
| | Horizondiscovery (Dharmacon) | siRNA KIAA1217/SKT 2 : 5'- GGAGGUGCAUGAUAUUGUU-3' | |
| | Horizondiscovery (Dharmacon) | siRNA KIAA1217/SKT 3 : 5'- GAACGCCUUUCUAAUGGAA-3' | |
| | Horizondiscovery (Dharmacon) | siRNA KIAA1217/SKT 4 : 5'- AACCAUCGAUUGCUUCUAA-3' | |
| p140Cap siRNA smart pool | Horizondiscovery (Dharmacon) | ref: M-010674-02-0005 | |
| MNR siRNA | Horizondiscovery (Dharmacon) | 5'-GCUCAAAGCUGAAGAAAUG-3' | Chevrier et al., 2016 |
| Scrambled siRNA | Horizondiscovery (Dharmacon) | ref: D-001206-14-05 | |
